# Crypt and Villus Enterochromaffin Cells are Distinct Stress Sensors in the Gut

**DOI:** 10.1101/2024.02.06.579180

**Authors:** Kouki K. Touhara, Nathan D. Rossen, Fei Deng, Tifany Chu, Andrea M. Harrington, Sonia Garcia Caraballo, Mariana Brizuela, Tracey O’Donnell, Onur Cil, Stuart M. Brierley, Yulong Li, David Julius

## Abstract

The crypt-villus structure of the small intestine serves as an essential protective barrier, with its integrity monitored by the gut’s sensory system. Enterochromaffin (EC) cells, which are rare sensory epithelial cells that release serotonin (5-HT), surveil the mucosal environment and signal both within and outside the gut. However, it remains unclear whether EC cells in intestinal crypts and villi respond to different stimuli and elicit distinct responses. In this study, we introduce a new reporter mouse model to observe the release and propagation of serotonin in live intestines. Using this system, we show that crypt EC cells exhibit two modes of serotonin release: transient receptor potential A1 (TRPA1)-dependent tonic serotonin release that controls basal ionic secretion, and irritant-evoked serotonin release that activates gut sensory neurons. Furthermore, we find that a thick protective mucus layer prevents TRPA1 receptors on crypt EC cells from responding to luminal irritants such as reactive electrophiles; if this mucus layer is compromised, then crypt EC cells become susceptible to activation by luminal irritants. On the other hand, villus EC cells detect oxidative stress through TRPM2 channels and co-release serotonin and ATP to activate nearby gut sensory fibers. Our work highlights the physiological importance of intestinal architecture and differential TRP channel expression in sensing noxious stimuli that elicit nausea and/or pain sensations in the gut.

## Introduction

Our gastrointestinal (GI) tract is equipped with a complex sensory system that detects the state of the gut mucosa and transmits signals within and outside this visceral organ. The first line of stimulus detection is mediated by enteroendocrine cells, which are rare specialized sensory cells within the gut epithelium that release hormones and neurotransmitters in response to endogenous and exogenous stimuli (*1*). Enterochromaffin (EC) cells are a subclass of excitable enteroendocrine cells that release serotonin (5-HT) in response to bacterial metabolites, neurotransmitters, interleukins, and ingested or endogenous irritants (*2-7*). Interestingly, EC cells also show spontaneous (basal) activity for which an underlying mechanism or physiologic role remains unknown (*6, 8*).

Several serotonin receptor subtypes are present in the gut (*5*), including ionotropic 5-HT3 and metabotropic 5-HT4 receptors, which have been pharmacologically targeted to treat GI dysregulation associated with diarrhea or constipation (*5, 9*). EC cells transduce signals to afferent sensory nerve fibers within the mucosa that express 5-HT3 receptors, excitatory ion channels that are activated by relatively high (micromolar) concentrations of serotonin (*4, 5, 10*). This serotonergic EC cell-sensory neuron circuit modulates a range of processes, including GI motility, secretion, nausea, and pain (*5, 11, 12*). In contrast, G protein-coupled 5-HT4 receptors on intestinal epithelial cells are activated by relatively low (nanomolar) levels of serotonin, leading to enhanced ionic secretion (*13, 14*). This latter process plays a crucial role in maintaining fluid balance in the gut, aiding digestion, and protecting the intestinal lining through the formation of a mucus barrier (*15*). However, it remains unclear if EC cells target both 5-HT3 and 5-HT4 receptors, or whether and when they exhibit the thousand-fold concentration difference in transmitter release that would be required to achieve receptor subtype-selective activation.

Such potential differential actions raise interesting questions about the temporal and spatial nature of serotonergic signaling in the intestine and how this relates to the arrangement of crypts and villi that define the complex architecture of the gut. Crypts are small invaginations within the epithelium that house stem cells, which are vulnerable to microbes and irritants and thus protected by antimicrobial peptides and a thick layer of mucus (*16*). Villi are long finger-like projections that extend from crypts towards the lumen and are more directly exposed to the luminal contents. Interestingly, EC cells change their molecular identity as they migrate from crypts to villi such that transient receptor potential A1 (TRPA1) channels are located predominantly in crypts, whereas TRPM2 channels are found mostly in villi (*17, 18*). However, it remains unclear whether mucus-protected EC cells in crypts or more exposed EC cells in villi respond to distinct stimuli to elicit differential physiological responses. Furthermore, the distribution of 5-HT receptors within the crypt-villus architecture, and how this relates to EC cell distribution, has not been elucidated. Addressing these important questions requires the development of new approaches for analyzing signaling in a complex organ structure with spatial and temporal resolution.

In this study, we describe a mouse model that allows for direct observation of serotonin release and propagation within the intact crypt-villus architecture. Using this system, we determine how differential release of serotonin from EC cells promotes distinct physiological responses within and beyond the gut, and how crypt and villus EC cells employ different TRP channels to detect exogenous or endogenous stress signals.

## Results

### Spatial and temporal dynamics of serotonin propagation

To monitor the real-time release and propagation of serotonin within the crypt-villus architecture, we set out to develop a mouse model that expresses genetically encoded GPCR-activation-based (gGRAB) serotonin sensors in the intestinal epithelium (*19, 20*). gGRAB_5HT3.0_ is an improved sensor whose fluorescence intensity increases upon serotonin binding (Figure 1A) (*20*). We developed a transgenic mouse line that expresses gGRAB_5HT3.0_ and a red-shifted Ca^2+^ indicator, jRGECO1a, following exposure to Cre recombinase (Figure S1A). Crossing the reporter line to Vil1-Cre mice resulted in the expression of gGRAB_5HT3.0_ and jRGECO1a in intestinal epithelial cells (Figure S1B), enabling us to visualize the release and propagation of serotonin within the gut using fluorescence microscopy (Figure 1B). It should be noted that the expression levels of gGRAB_5HT3.0_ and jRGECO1a decrease progressively in the proximal and distal colon, consistent with the expression pattern of the Villin gene (*21*) (Figure S1B). To visualize reporter activation, we removed a section of jejunum, flushed the luminal contents, and then filleted the tissue to create a flat sheet. We imaged the tissue either from the smooth muscle side to visualize crypts or from the luminal side to observe villi. When exposed to a high K^+^ solution, EC cells released serotonin, which subsequently activated the gGRAB_5HT3.0_ sensor in both EC cells and adjacent epithelial cells within crypts and villi (Figure 1C and 1D, and Supplementary videos 1 and 2). Serendipitously, we observed the highest fluorescence intensity of gGRAB_5HT3.0_ in EC cells in either fixed or live tissues, which facilitated the identification of EC cells during imaging (Figure 1E-1H).

**Figure 1.**
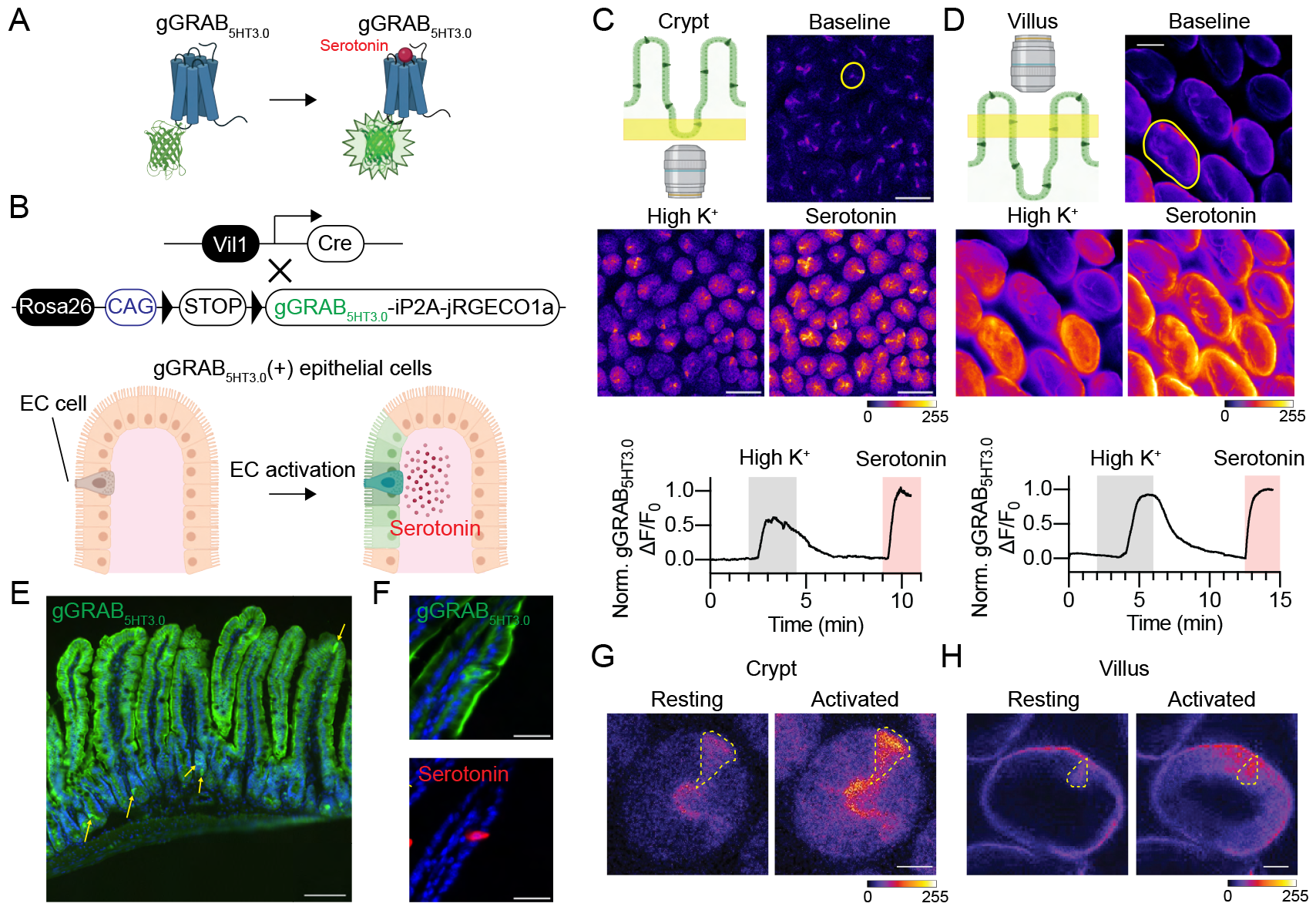
Live imaging of the EC-derived serotonin in the intact crypt-villus architecture. (A) The gGRAB_5HT3.0_ sensor is a modified metabotropic 5-HT receptor fused to cpEGFP. Upon serotonin binding, the receptor changes its conformation, leading to an increase in the fluorescent intensity of cpEGFP. (B) Generation of Vil1-Cre;gGRAB_5HT3.0_ mice that express gGRAB_5HT3.0_ sensors and jRGECO1a in intestinal epithelial cells, allowing for visualization of the release and propagation of EC-derived serotonin. (C) and (D) Examples of gGRAB_5HT3.0_ sensor imaging in crypts (C) and villi (D). Serotonin release was stimulated by application of high K^+^ (70 mM KCl). The gGRAB_5HT3.0_ sensor was fully activated with 20 µM serotonin at the end of recordings for normalization. Yellow circles indicate the region of interest (ROI) used for graph plotting. Scale bar = 100 µm. (E) Expression pattern of gGRAB_5HT3.0_ sensors in the small intestine. The gGRAB_5HT3.0_ sensor was immunostained with an anti-GFP antibody. Arrows point to the EC cells in crypts and villi. Scale bar = 100 µm. (F) The immunofluorescence intensity of the gGRAB_5HT3.0_ sensor is higher in EC cells compared to other epithelial cells. Scale bar = 30 µm. (G) and (H) Example of live gGRAB_5HT3.0_ sensor imaging in crypt EC (G) and villus EC (H) cells following high K^+^ activation. EC cells are outlined as dashed yellow lines. Scale bar = 20 µm.

### Two modes of TRPA1-dependent serotonin release

Having established a tool to visualize serotonin release in the intact intestine, we observed an interesting differential profile whereby tonic serotonin release was seen in crypts but not villi (Figure 2A and Supplementary video 3). This tonic release was similarly observed when crypts were isolated from intact tissue (Figure S2A). We noticed that this differential activity aligned with preferential expression of TRPA1 (the ‘wasabi receptor’) in crypt EC cells (17, 18). TRPA1 is a chemo-nociceptive ion channel that functions as a detector of chemically reactive electrophiles, including plant-derived irritants, environmental toxicants, and endogenous products of oxidative stress. Indeed, *in situ* hybridization histochemistry revealed robust *Trpa1* expression in crypts that diminished progressively from lower to upper villi (Figure S2B). In addition, we observed serotonin release from dissociated crypts, but not from similarly dissociated upper villi, upon stimulation with the TRPA1 agonist, allyl isothiocyanate (AITC) (Figure S2C). This observation confirms the functional expression of TRPA1 channels in crypts but not in upper villi. To determine whether TRPA1 contributes to tonic serotonin release in crypt EC cells, we used intestinal organoids derived from Tac1-Cre;GCaMP5g-IRES-tdTomato mice, which express the Ca^2+^ indicator GCaMP5g and tdTomato specifically in EC cells (Figure S2D). These organoids replicate crypt features, thereby representing crypt EC cells (*17*). In this system, a TRPA1 antagonist, A967079 (A96), diminished the spontaneous activity of EC cells (Figure 2B). Consistently, we observed A96-sensitive spontaneous TRPA1 channel openings in single-cell voltage-clamp recordings from EC cells (Figure S2E) and found that A96-sensitive spontaneous membrane depolarizations drove repeated action potentials (Figure S2F). Additionally, spontaneous calcium flux in EC cells was inhibited by tetrodotoxin (TTX), a voltage-gated sodium channel (NaV) blocker (Figure S2G). Taken together, these results suggest that low levels of TRPA1 channel opening are sufficient to drive NaV-dependent action potentials, leading to tonic serotonin release from crypt EC cells.

**Figure 2.**
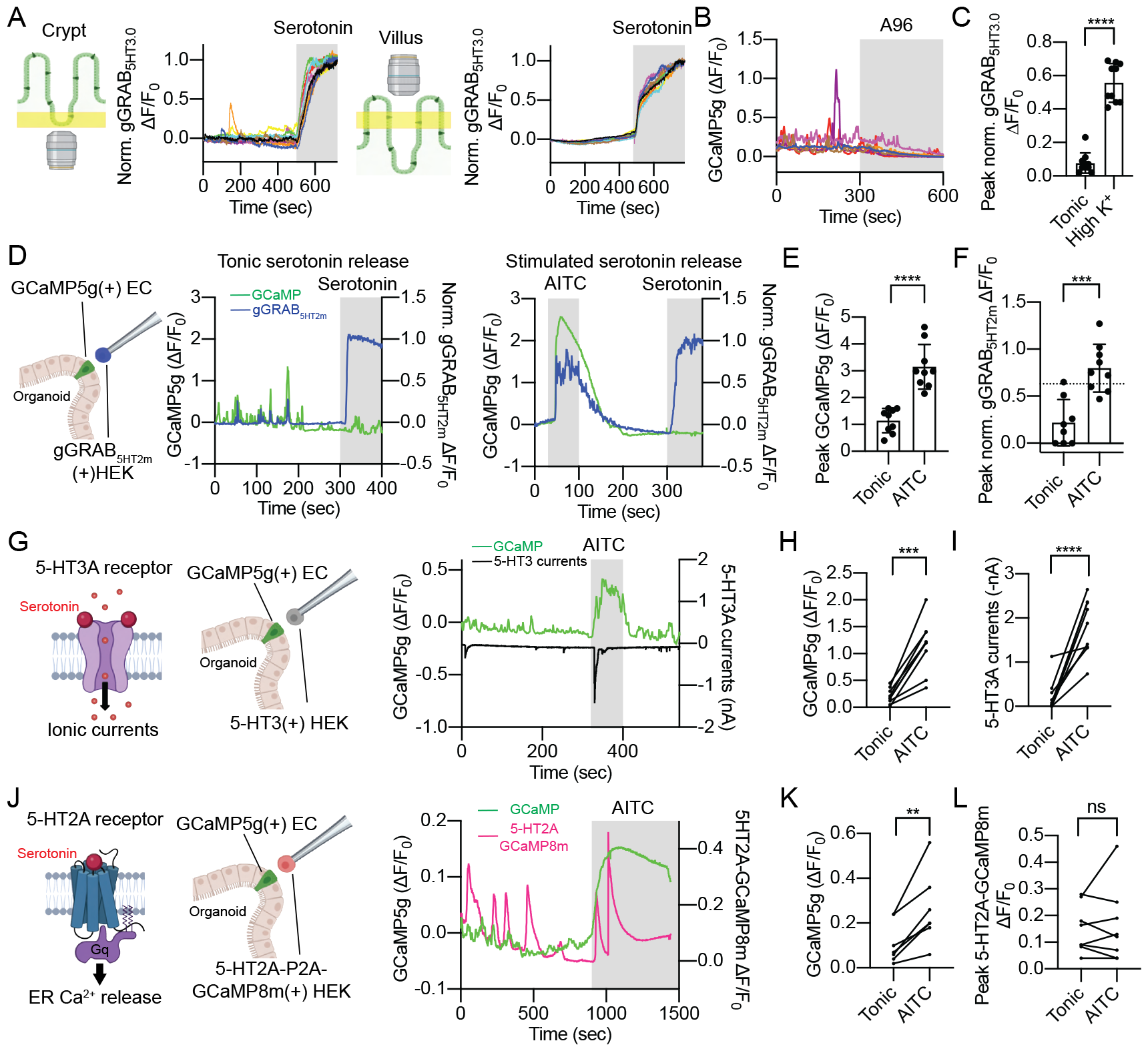
Crypt EC cells show two modes of serotonin release dependent on TRPA1. (**A**) Basal serotonin release in crypts (left) or villi (right). gGRAB_5HT3.0_ was fully activated with 20 µM serotonin at the end of recordings for normalization. (B) 10 µM A96, a TRPA1 antagonist, inhibits the tonic activity of crypt EC cells in intestinal organoids. (C) Peak normalized gGRAB_5HT3.0_ response in crypts during the tonic and high K^+^ stimulated phases. Welch’s t-test; ****p ≤ 0.0001; n = 10. (D) In each experiment, a HEK cell expressing the gGRAB_5HT2m_ sensor was positioned 5 µm away from a GCaMP5g-expressing organoid EC cell. The gGRAB_5HT2m_ and GCaMP5g signals were simultaneously measured during tonic or AITC-stimulated activity. gGRAB_5HT2m_ was fully activated with 500 µM serotonin at the end of recordings for normalization. (E) Peak GCaMP5g response of EC cells during tonic vs. AITC-stimulated activity. Welch’s t-test; ****p ≤ 0.0001; n = 8 and 9. (F) Peak normalized gGRAB_5HT2m_ response during tonic vs. AITC-stimulated activity. The dashed line indicates a normalized gGRAB_5HT2m_ response of 0.63, corresponding to 1 µM serotonin. Welch’s t-test; ***p ≤ 0.001; n = 8 and 9. (G) Single-cell patch-clamp recordings were performed on HEK cells positioned 5 µm away from GCaMP5g-expressing EC cells in organoids. Inward 5-HT3 currents were measured while the GCaMP5g signal was recorded. (H) Peak GCaMP5g response of EC cells during the tonic and AITC-stimulated phases. Paired t-test; ***p ≤ 0.001; n = 9. (I) Maximum inward 5-HT3 currents during tonic vs. AITC-stimulated activity. Paired t-test; ****p ≤ 0.0001; n = 9. (J) In each experiment, a HEK cell expressing GCaMP8m and 5-HT2A receptors was positioned 5 µm away from a GCaMP5g-expressing organoid EC cell. GCaMP signals from HEK and EC were simultaneously measured. (K) Peak GCaMP5g response of EC cells during the tonic and AITC-stimulated phases. Paired t-test; **p ≤ 0.01; n = 8. (L) Peak 5-HT2A-GCaMP8m biosensor response during tonic vs. AITC-stimulated activity. Paired t-test; ns: not significant; n = 8.

The normalized gGRAB_5HT3.0_ sensor response suggested that tonic serotonin release is of smaller magnitude compared to stimulated release (Figure 2C). To verify this observation, we quantitatively compared tonic and AITC-stimulated serotonin release from crypt EC cells in intestinal organoids. For this experiment, we used the gGRAB_5HT2m_ sensor, whose affinity is more suitable for measuring serotonin within the micromolar range (*20*). We positioned a human embryonic kidney (HEK293) cell expressing the gGRAB_5HT2m_ sensor adjacent to (5 μm away from) an EC cell in Tac1-Cre;GCaMP5g-IRES-tdTomato organoids. While observing the spontaneous and AITC-stimulated GCaMP signals in EC cells within organoids, we simultaneously monitored the gGRAB_5HT2m_ sensor signal in the juxtaposed biosensor cell (Figure 2D). At the end of each recording, we applied a maximally effective concentration of serotonin to fully activate the gGRAB_5HT2m_ sensor. This normalization step enabled us to estimate the local concentration of released serotonin, based on the dose-response curve of the gGRAB_5HT2m_ sensor (Figure S2H). We observed a lower GCaMP signal during the tonic phase compared to the more robust signal observed during the AITC-stimulated phase (Figure 2E). Furthermore, we found that the amount of serotonin released during the tonic phase was significantly lower than that elicited in response to AITC-evoked TRPA1 activation (Figure 2F). Specifically, tonic serotonin release remained at a minimal level: the normalized gGRAB_5HT2m_ amplitude surpassed 0.63 (corresponding to 1 μM serotonin) in just 1 of 8 cells examined. Conversely, for AITC-stimulated release, the normalized gGRAB_5HT2m_ amplitude exceeded 0.63 in 6 of 9 cells evaluated.

If tonic serotonin release is in the nanomolar range (and distinctively lower than AITC-evoked release in the micromolar range), then tonic serotonin should predominantly activate metabotropic 5-HT receptors, but not ionotropic 5-HT3 receptors, whereas stimulated serotonin should activate both receptor types. To test this hypothesis, we first monitored GCaMP signals in EC cells within organoids to detect their activation while simultaneously measuring whole-cell currents in neighboring HEK293 biosensor cells expressing the ionotropic 5-HT3 receptor, which is activated by serotonin in the micromolar range (Figure 2G) (*10*). We found that peak 5-HT3 currents were substantially smaller during tonic serotonin release compared to those observed during AITC-stimulated release, consistent with the idea that EC cells activate ionotropic 5-HT3 receptors most robustly when stimulated by agonists (Figure 2G-2I).

On the other hand, tonic serotonin release should be sufficient to activate high-affinity metabotropic 5-HT receptor subtypes such as 5-HT4 and 5-HT2, both exhibiting nanomolar sensitivity to serotonin (*13, 22*). To test this prediction, we developed a biosensor in which HEK293 cells co-express Gq-coupled 5-HT2A receptors and GCaMP8m. When activated, 5-HT2A receptors promote endoplasmic reticulum (ER)-stored Ca^2+^ release, which results in increased GCaMP8m fluorescence. Notably, low-nanomolar concentrations of serotonin repeatedly activated this biosensor without apparent desensitization (Figure S2I). Despite observing a more robust response in EC cells during the AITC-stimulated phase, comparisons of peak biosensor responses during tonic and AITC-stimulated release revealed that both phases activated the biosensor to similar extents (Figure 2J-2L). Taken together, the gGRAB_5HT2m_, 5-HT3, and 5-HT2A biosensor experiments are in complete agreement and suggest that tonic nanomolar serotonin activates metabotropic rather than ionotropic 5-HT receptors. Conversely, when crypt EC cells are stimulated by electrophiles (or other agonists), they release micromolar concentrations of serotonin, activating both metabotropic and ionotropic 5-HT receptors.

### Distribution of 5-HT receptors in the crypt-villus structure

What are the physiological consequences of tonic (nanomolar) versus stimulated (micromolar) serotonin release in crypts? We focused on 5-HT3 and 5-HT4 subtypes as the most physiologically relevant targets in the intestine (*5, 9*). We first investigated the distribution of the 5-HT4 receptor, which is known to stimulate epithelial chloride secretion, thereby influencing the rate of fluid transfer into the intestinal lumen (*14*). Interestingly, *in situ* hybridization histochemistry revealed that 5-HT4 receptors are localized in crypts, but not villi (Figure 3A and 3B). This distribution, and low-nanomolar affinity for serotonin, suggest a potential preference for activation when serotonin is released tonically from crypt EC cells. To test this hypothesis, we used an Ussing chamber to measure ionic secretion. Consistent with previous findings, serotonin stimulated ionic secretion in a 5-HT4-dependent manner (Figure 3C and S3A) (*14*). To specifically activate EC cells in this system, we used Tac1-Cre;ePet-Flp;hM3Dq mice, in which deschloroclozapine (DCZ) triggers serotonin release from EC cells expressing the excitatory DREADD receptor hM3Dq (*12*). Application of DCZ indeed stimulated ionic secretion, revealing a role for EC cells in 5-HT4-mediated fluid secretion (Figure 3D and S3A). We next investigated the contribution of tonic serotonin release to basal ionic secretion using intestinal organoids. Activation of the stimulatory G protein (G_s_) pathway is known to induce fluid secretion, leading to swelling of organoids (*23*). Indeed, acute exposure to serotonin produced organoid swelling in a 5-HT4-dependent manner (Figure 3E). Furthermore, a reduction in basal swelling was observed when organoids were incubated with a TRPA1 antagonist (A96) or a 5-HT4 antagonist (RS) (Figure 3F), suggesting that TRPA1-induced tonic serotonin release contributes to organoid swelling via activation of 5-HT4 receptors. Notably, long-term exposure to A96 and RS did not affect overall organoid growth, excluding the potential impact of these drugs on organoid proliferation (Figure S3B). Based on these findings, we conclude that tonic serotonin release from crypt-residing EC cells activates 5-HT4 receptors, predominantly localized in crypts, thereby stimulating ionic secretion.

**Figure 3.**
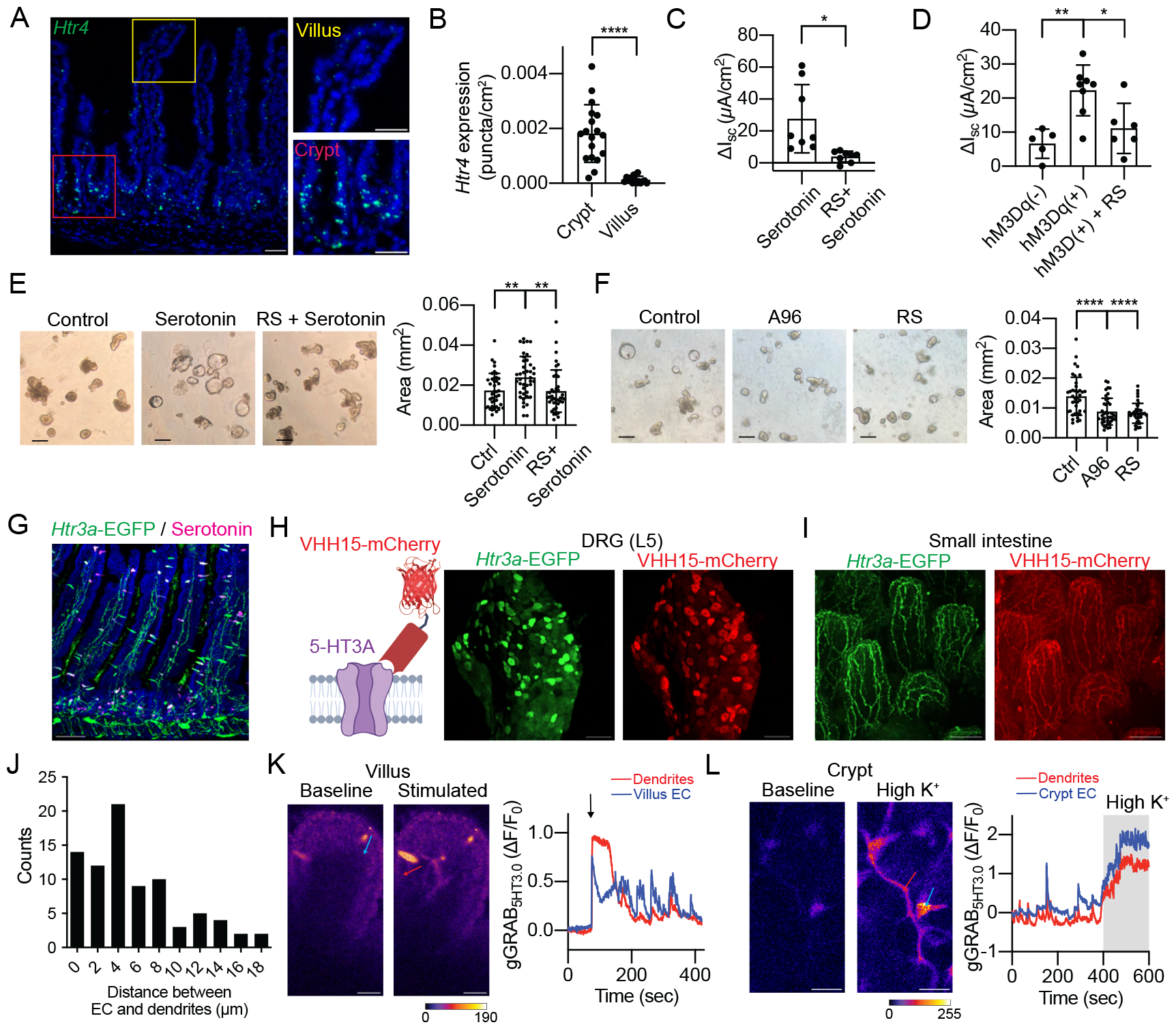
Metabotropic and ionotropic 5-HT receptors reside in proximity to crypt EC cells. (A) *In situ* hybridization of 5-HT4 receptors (*Htr4*) in the small intestine. Magnified images of villus (yellow) and crypt (red). Scale bar = 20 µm. (B) Quantification of *in situ* hybridization of *Htr4* in crypts and villi. Welch’s t-test; ****p ≤ 0.0001; n = 15 and 18. (C) Summary of changes in Isc (ΔIsc) induced by 1 µM serotonin with or without pretreatment with a 5-HT4 antagonist (1 µM RS 23597-190). Welch’s t-test; *p ≤ 0.05; n = 8. (D) Summary of changes in Isc (ΔIsc) induced by1 µM DCZ in the ileal mucosa isolated from Tac1-Cre;ePet-Flp;hM3Dq mice, with or without 1 µM RS 23597-190. These mice express hM3Dq DREADD receptors specifically in EC cells. ePet-Flp negative litter mates were used as control animals. Ordinary one-way ANOVA; **p ≤ 0.01; *p ≤ 0.05; n = 5-8. (E) Day 1 intestinal organoids were incubated with 1 µM serotonin with or without 10 µM RS 23597-190 for 60 minutes. Organoids exhibited 5-HT4-dependent swelling (left). Cross-sectional area of organoids was compared among untreated (ctrl), serotonin-treated, and RS + serotonin-treated conditions (right). n = 40-41. Ordinary one-way ANOVA; **p ≤ 0.01. Scale bar = 50 µm. (F) Day 1 intestinal organoids were incubated with 5 µM A96 or 10 µM RS for 12 hours, and cross-sectional areas were compared. n = 40. Ordinary one-way ANOVA; ****p ≤ 0.0001. Scale bar = 50 µm. (G) *Htr3a*-EGFP mice express cytosolic EGFP in the nerve fibers (Green). Serotonin immunofluorescence is shown in magenta. *Htr3a*-EGFP mice also show ectopic expression in epithelial cells. Scale bar = 100 µm. (H) Immunohistochemical labeling of 5-HT3 receptors using a specific 5-HT3A nanobody conjugated to mCherry (VHH15-mCherry). Dorsal root ganglion (L5) isolated from *Htr3a*-EGFP mice (green) were labeled with VHH15-mCherry (red). Signals were amplified by immunostaining for EGFP and mCherry. Scale bar = 100 µm. (I) Immunohistochemical labeling of 5-HT3 receptors in nerve fibers. The small intestine isolated from *Htr3a*-EGFP mice (green) was labeled with VHH15-mCherry (red). Signals were amplified by immunostaining for EGFP and mCherry. Scale bar = 100 µm. (J) Distribution of the distances (in µm) between the basolateral side of EC cells and the nearest nerve fibers. Average = 5.5 ± 4.5 µm; Median = 4.7 µm (n = 82, ± SD). (K) Simultaneous gGRAB_5HT3.0_ imaging of villus EC cells (blue arrow) and nerve fibers (red arrow) using Insm1-Cre;gGRAB_5HT3.0_ mice. Villus EC cells were electrically stimulated as indicated above the signal. Scale bar = 30 µm. (L) Simultaneous gGRAB_5HT3.0_ imaging of crypt EC cells and mucosal nerve fibers using Insm1-Cre;gGRAB_5HT3.0_ mice. Crypts EC cells were stimulated by high K^+^ (70 mM KCl) at the end of the recording. Scale bar = 30 µm.

We next asked whether crypt and villus EC cells can target ionotropic 5-HT3 receptors, which require micromolar serotonin for activation. It is known that both intrinsic and extrinsic sensory afferents express ionotropic 5-HT3 receptors (*12, 24, 25*). In a reporter mouse expressing cytosolic GFP under the control of the 5-HT3A promoter, nerve fibers innervating the villi are labeled with GFP (Figure 3G). Yet, this reporter line does not precisely pinpoint the localization of 5-HT3 receptors along the nerve fibers. Therefore, it remained unclear if 5-HT3 receptors are preferentially localized near certain EC cells or broadly distributed along the nerve fibers. To visualize 5-HT3 receptors themselves, we deployed a modified 5-HT3A-specific nanobody (VHH15) developed previously for structure determination (Figure 3H) (*26*). A VHH15-mCherry fusion specifically labeled 5-HT3 receptors ectopically expressed in HEK293 cells, as well as native 5-HT3 receptors in dorsal root ganglia (Figure 3H and S3C). Moreover, VHH15-mCherry labeled nerve fibers innervating the mucosa in both the small and large intestine, revealing uniform expression of these channels along afferents (Figure 3I and S3D).

Our serotonin staining therefore implies that EC cells, whether located in crypts or villi, can activate sensory neurons innervating the mucosa if the released transmitter reaches them at a sufficiently high (micromolar) concentration required for 5-HT3 receptor activation. To directly assess the propagation of EC-derived serotonin, we first established the distance between the basolateral side of EC cells and the closest nerve fibers (Figure 3J and S3E). This distance (average = 5.5 ± 4.5 μm) is comparable to that between EC and biosensor cells used in quantification experiments described above, indicating that most EC cells can present micromolar levels of serotonin to mucosal afferents when stimulated. Furthermore, using Insm1-Cre;gGRAB_5HT3.0_ sensor mice, which express gGRAB_5HT3.0_ in both EC cells and nerve fibers (Figure S3F), we could directly observe the propagation of serotonin between these entities. Whether released from crypt or villus EC cells, serotonin readily reached and covered nearby nerve fibers (Figure 3K-3L and S3G-S3H). In summary, our immunohistochemical and gGRAB_5HT3.0_ sensor analyses show that both crypt and villus EC cells are sufficiently close to 5-HT3 receptors on mucosal sensory nerve fibers to transmit excitatory serotonergic signals to these afferents.

### Loss of protective mucosal layer unmasks TRPA1 sensitivity

What physiologic circumstances might promote bolus serotonin release from crypt EC cells? As noted above, TRPA1 is an irritant-activated ion channel that responds to a wide range of electrophilic toxicants and inflammatory agents (*27, 28*). These include pungent agents from wasabi, garlic, onion, and other members of the *Brassica* and *Allium* plant family, which are associated with both potential health benefits and risks (*29, 30*). Also, environmental toxicants or metabolic byproducts of certain chemotherapeutic drugs are strong electrophiles that elicit severe inflammation in internal organs (*31*). We were therefore curious to know whether such dietary electrophiles (AITC, allicin, and cinnamaldehyde) or metabolites (acrolein and 2-pentenal) could activate crypt EC cells. In organoids from Tac1-Cre;GCaMP5g-IRES-tdTomato mice, each of these agents elicited robust responses in EC cells (Figure 4A and S4A) that were blocked by the TRPA1 antagonist, A96 (Figure 4A and S4A).

**Figure 4.**
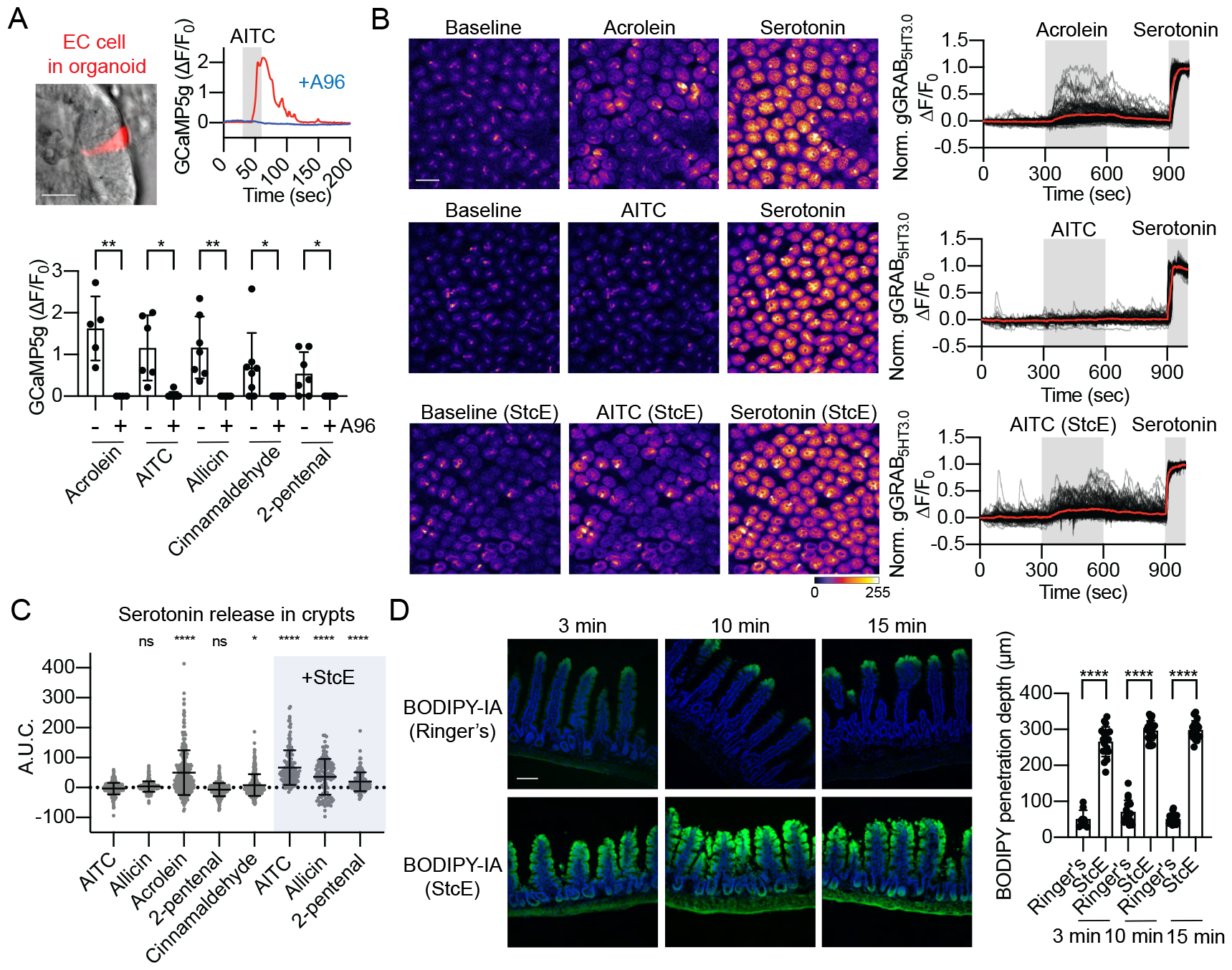
TRPA1 is an electrophile sensor in the mucus-protected crypt EC cell. (A) A wide range of electrophiles activate EC cells within intestinal organoids derived from Tac1-Cre;GCaMP5g-IRES-tdTomato mice. EC cells were stimulated by indicated electrophiles (100 µM) in the presence or absence of 100 µM A96. Scale bar = 20 µm. Welch’s t-test; *p ≤ 0.05; **p ≤ 0.01; n = 4-8. (B) and (C) Electrophile activation of crypt EC cells *ex vivo*. Jejunal tissue was pretreated with StcE or saline and subsequently exposed to the indicated electrophiles (100 µM) for 5 minutes. gGRAB_5HT3.0_ sensors were fully activated with 20 µM serotonin at the end of the recording, and signals were normalized to the fully activated value. The area under the curve (A.U.C.) was calculated during the agonist application (300-600 sec). Scale bar = 30 µm. Ordinary one-way ANOVA; *p ≤ 0.05; ****p ≤ 0.0001; ns: not significant. n = 138-278. (D) Mucus limits the diffusion of electrophiles into the mucosal tissue. The ex vivo jejunal prep was exposed to 10 mM StcE or saline for 60 minutes. The tissue was then stained with 10 µM BODIPY-IA for 3-15 minutes and the penetration depth was measured from the villus tips. Welch’s t-test; ****p ≤ 0.0001; n = 9-21. Scale bar = 100 µm.

We next examined the effects of electrophiles in freshly prepared gut tissue, where the crypt structure and its protective mucus layer are preserved. Using gGRAB_5HT3.0_ mice, we were surprised to find that only acrolein, a highly reactive electrophile, robustly stimulated crypt EC cells, whereas other electrophiles were ineffective (Figure 4B, 4C, and S4B). We reasoned that if the mucus layer acts as a barrier to irritant access, then its degradation should increase susceptibility of crypt EC cells to weaker electrophiles. To test this hypothesis, we incubated gut tissue with StcE, a mucinase from pathogenic *E*.*coli* O157 that digests Muc2, a primary component of intestinal mucus (*32-34*). To determine whether StcE digestion enhances crypt access, we exposed gut tissue to a fluorescently tagged electrophile, BODIPY-iodoacetamide (BODIPY-IA). In the presence of an intact mucus layer, BODIPY-IA infiltrated only the villus tips, even after 15 minutes of incubation (Figure 4D). Remarkably, StcE treatment allowed BODIPY-IA to reach the crypts in as little as 3 minutes, supporting the idea that the mucus layer restricts electrophile access to the crypt region (Figure 4D). Subsequently, we exposed StcE-treated intestinal tissue from gGRAB_5HT3.0_ mice to previously ineffective electrophiles (AITC, allicin, and 2-pentenal) and found that they significantly activated crypt EC cells after mucus digestion (Figure 4B, 4C, and S4C). We conclude from these experiments that TRPA1 channels on crypt EC cells can respond to electrophiles, but under normal conditions a protective mucus layer permits only very potent and permeable electrophiles, such as acrolein, to access EC cells and activate TRPA1. However, when this protective mucus barrier is degraded, other electrophilic irritants, such as those from dietary sources, can also gain access to crypt EC cells and activate TRPA1 to stimulate serotonin release.

### TRPM2 is an oxidative stress sensor in the villus

Given that TRPA1 is preferentially expressed by crypt EC cells (Figure S2B), we wondered if villus EC cells also have the capacity to detect chemical irritants or other indicators of tissue damage. Interestingly, our *in situ* hybridization analysis showed preferential expression of TRPM2 channels in villus EC cells (Figure 5A), again consistent with available single-cell RNA sequencing data (*18*). TRPM2 is activated by intracellular ADP-ribose, which is generated during oxidative stress (*35*). Indeed, whole-cell patch-clamp recordings revealed ADP-ribose-activated inward currents in EC cells (Figure 5B). These currents were attenuated by 2-aminoethoxydiphenyl borate (2-APB), which inhibits TRPM2, and reduced upon replacement of extracellular Na^+^ with NMDG^+^. Consistent with the expression pattern of TRPM2, we observed significantly larger ADP-ribose-activated currents in villus versus crypt EC cells (Figure 5B).

**Figure 5.**
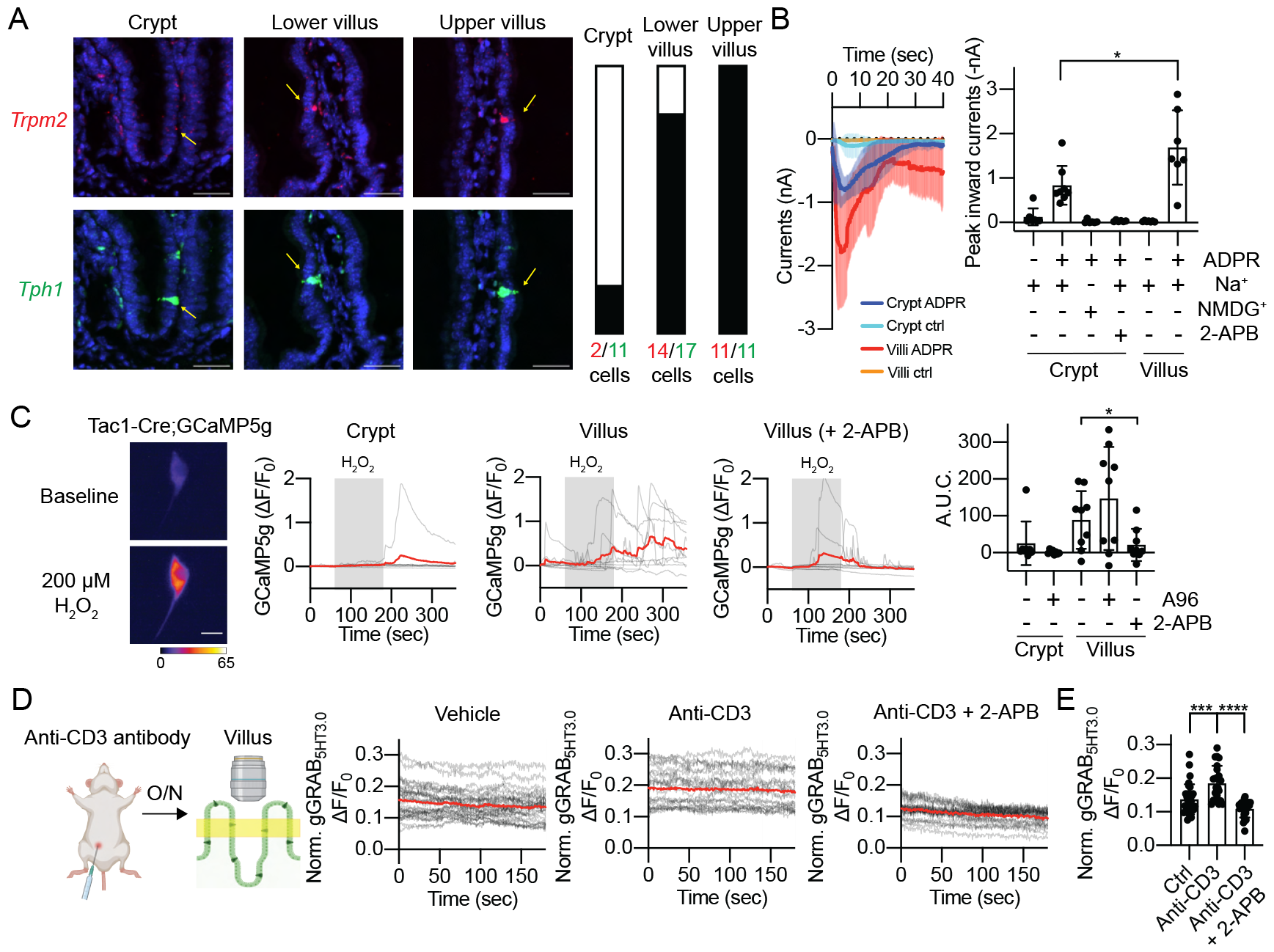
TRPM2 is an oxidative stress sensor in the villus EC cell. (A) *In situ* hybridization of *Trpm2* and *Tph1* in the small intestine. Yellow arrows indicate *Tph1*(+) EC cells. Bars indicate the ratio of *Trpm2*-positive cells / *Tph1*-positive cells. Scale bar = 30 µm. (B) Voltage-clamp recordings of ADP-ribose (ADPR)-induced TRPM2 currents (left). The recordings reached whole-cell configuration at time = 0 and membrane potential was held at -80 mV. The recordings were performed in Na^+^ or NDMG^+^ extracellular solution, in the absence or presence of a TRPM2 antagonist, 30 µM of 2-APB (right). Ordinary one-way ANOVA; *p ≤ 0.05. n = 6-8. (C) Oxidative stress activates villus EC cells. Dissociated EC cells expressing GCaMP5g were exposed to 200 µM H2O2 for 2 minutes in the presence or absence of 100 µM A96 or 30 µM 2-APB. Scale bar = 10 µm. Ordinary one-way ANOVA; *p ≤ 0.05. n = 8-9. (D) and (E) Anti-CD3 antibody-induced oxidative stress activates villus EC cells. The epithelial inflammation was induced by anti-CD3 antibody in gGRAB_5HT3.0_ sensor mice, and basal serotonin levels were measured in fleshly isolated *ex vivo* prep in the presence or absence of 60 µM 2-APB. Ordinary one-way ANOVA; ***p ≤ 0.001. n = 23-31.

We next asked whether oxidative stress activates villus EC cells. Using dissociated EC cells expressing GCaMP5g, we found that 200 μM H_2_O_2_ robustly activated villus EC cells in a TRPM2-(but not TRPA1) dependent manner (Figure 5C and S5A). Furthermore, in a small intestinal injury model induced by administration of an anti-CD3 antibody (*36*), we observed a TRPM2-dependent increase in serotonin release in villi but not in crypts (Figure 5D-5E and S5B), suggesting that TRPM2 channels sense inflammatory stress of epithelial cells within villi.

### Villus EC cells co-release serotonin and ATP

Villus EC cells do not exhibit tonic serotonin release (Figure 2A), and the expression of 5-HT4 receptors in villi is minimal (Figure 3A and 3B). Given these findings, we hypothesized that villus EC cells might have distinct neurotransmitter release patterns and targets. While it has been suggested that EC cells release ATP alongside serotonin (*37*), this has not been thoroughly investigated. To explore this, we performed a biosensor experiment with a genetically encoded ATP sensor, gGRAB_ATP1.0_ (*38*). While observing high K^+^-stimulated GCaMP signals in EC cells within organoids or primary isolated EC cells, we simultaneously monitored the gGRAB_ATP1.0_ sensor signal in neighboring biosensor cells (Figure 6A). We generated villus-like organoids by treating them with BMP4, a key regulator of crypt-villus axis differentiation (*17*). Interestingly, we found that EC cells released ATP in villus-differentiated organoids and primary dissociated villus cells, but not in organoids where crypt EC cells predominate or in isolated primary crypts (Figure 6A and 6B). Moreover, release was attenuated in the absence of extracellular Ca^2+^, suggesting that ATP is co-released with serotonin *via* secretory vesicles (Figure 6B). To ask whether released nucleotide contributes to sensory nerve fiber activation, we next investigated the sensitivity of mucosal afferents to ATP and serotonin. We traced and dissociated mucosa-innervating vagal neurons from the small intestine and performed single-cell Ca^2+^ imaging (Figure 6C) (*39*). Remarkably, all traced mucosal vagal neurons responded to ATP, and a majority also responded to the 5-HT3 receptor-selective agonist, mCPBG (Figure 6D and 6E). Consistent with these results, single-cell real-time PCR from these neurons revealed the expression of P2X and 5-HT3 receptors (Figure S5C), demonstrating that villus EC-derived ATP and serotonin collaborate to activate mucosal vagal afferents. Moreover, most of these neurons did not respond to AITC or H_2_O_2_, or express TRPA1 or TRPM2 channels, emphasizing the role of EC cells as specialized sensors for electrophiles and oxidative stress in the small intestine that couple to mucosal afferents (Figure 6D-6E and S5C).

**Figure 6.**
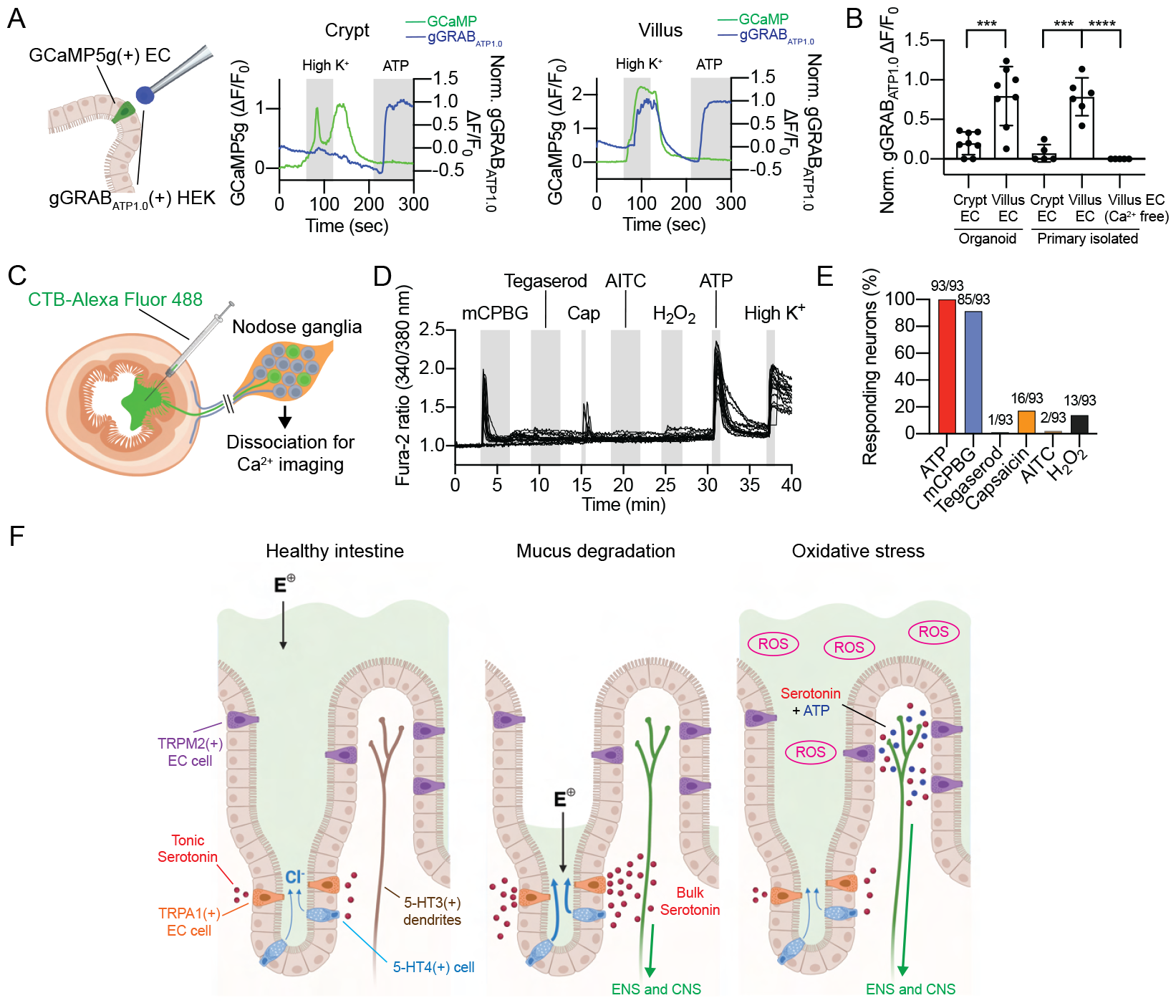
Villus EC cells co-release serotonin and ATP. (A) and (B) Villus but not crypt EC cells release ATP. In each experiment, a HEK cell expressing the gGRABATP1.0 sensor was positioned 5 µm away from a GCaMP5g-expressing EC cell. The signals from the gGRABATP1.0 sensor and GCaMP5g were simultaneously measured from the crypt and villus-differentiated organoids, and primary isolated crypts and villi. EC cells were stimulated by high K^+^ (70 mM KCl). The gGRABATP1.0 sensor was fully activated with 20 µM ATP at the end of recordings for normalization. Welch’s t-test; **p ≤ 0.01; ***p ≤ 0.001; n = 5-9. (C) Green fluorescent-conjugated cholera toxin B (CTB) retrograde tracer was injected into the small intestine lumen to specifically trace mucosal afferents while excluding fibers innervating the muscle layer. The retrogradely traced vagal neurons were dissociated for calcium imaging and single-cell real-time PCR analysis. (D) Representative calcium imaging traces of dissociated small intestine mucosa-innervating vagal neurons. 10 µM mCPBG (5-HT3 agonist), 1 µM tegaserod (5-HT4 agonist), 50 nM capsaicin (TRPV1 agonist), 1 µM AITC, 200 µM H2O2, 10 µM ATP, and 40 mM KCl (high K^+^) were applied as indicated above the signals. (E) The percentage of small intestine mucosa-innervating vagal neurons responding to the applied agonists. n = 93 neurons. (F) In the healthy intestine, a layer of mucus shields the epithelium from luminal electrophiles (E+), while the tonic, TRPA1-dependent activity of EC cells regulates ionic secretion from crypts. Upon damage to the mucus layer, luminal electrophiles further activate crypt EC cells, stimulating a bolus of serotonin release that activates the mucosal nerve fibers to relay signals to the enteric and central nervous systems (ENS and CNS). On the other hand, villus EC cells are silent in the healthy intestine. These cells detect elevated reactive oxygen species (ROS) via activation of TRPM2 and respond by co-releasing serotonin and ATP.

## Discussion

### Two modes of serotonin release

In this study, we develop and exploit biosensors to characterize spatial and temporal dynamics of neurotransmitter signaling within the gut architecture with the goal of understanding the relevance of these parameters to homeostatic and protective functions. Previously, high-performance liquid chromatography (HPLC) and enzyme-linked immunosorbent assay (ELISA) were employed to measure gut serotonin (*40, 41*). However, these methods primarily measure serotonin extracted from entire tissue samples and typically provide single timepoint measurements without spatial information. A more recent innovation, the tissue-like electrochemical biosensor, was designed to record serotonin in the intact intestine, but this method predominantly measures luminal serotonin (*42*). By comparison, our approach provides enhanced spatial, temporal, and quantitative analyses that greatly enhance our understanding of how specific stimuli trigger transmitter release from sensory cells within a complex anatomical structure - in this case, crypt-villus architecture with an intact mucus layer. By directing expression of gGRAB_5HT3.0_ sensors to all intestinal epithelial cells, we can visualize the extent of diffusion of EC-derived serotonin within the epithelial layer following activation. Another noteworthy feature of the gGRAB_5HT3.0_ sensor is its independence from arrestin-mediated desensitization (*20*). This allows for extended (> 30 minutes) imaging, as well as signal normalization upon addition of a saturating concentration of agonist, thereby increasing the method’s quantitative robustness.

Using these tools, we found that EC cells within the crypt release serotonin in two ways, including low-level tonic and high-level evoked modes. One interesting question is what accounts for TRPA1-dependent tonic release? Our patch-clamp experiments show that basal TRPA1 activity can be observed in cultured EC cells and that single-channel events depolarize the membrane sufficiently to activate NaV channels and elicit action potentials. Thus, tonic transmitter release is likely a cell autonomous process driven by low-level TRPA1 activity that is either spontaneous or elicited by tonic low-level production of cellular electrophiles. TRPA1 is also a ‘receptor operated’ channel that can be activated downstream of signaling pathways that increase cytosolic calcium, representing another potential regulatory mechanism (*43*). The identification of factors or conditions that support tonic channel activation may provide insights into endogenous or metabolic processes that regulate basal EC cell excitability. In any case, our findings suggest that crypt EC cells constitutively modulate ionic secretion through 5-HT4 receptor-expressing cells located within the crypt. Consistent with this, it has been shown that a 5-HT4 antagonist decreases basal ionic secretion in the small intestine (*14*). Therefore, crypt EC cells may fine-tune ionic secretion in response to changes in luminal or endogenous electrophiles that access crypts. Importantly, this low level of tonically released serotonin does not activate 5-HT3 receptors on sensory neurons, suggesting that crypt EC cells control gut secretion without conveying signals to intrinsic or extrinsic sensory networks, except perhaps in extreme pathological circumstances that degrade the protective mucus layer (see below).

It has been proposed that enteroendocrine L cells communicate with sensory neurons through synapse-like contacts (*44*), an idea that we subsequently suggested might also apply to interactions between EC cells and primary afferents (*6*). While a subset of EC cells and nerve fibers are closely juxtaposed, our current analysis indicates that the majority are too distant to establish *bona fide* synapses and likely communicate in a paracrine manner. Specifically, we show that released serotonin diffuses towards the closest dendrites, which uniformly express 5-HT3 receptors along their length. Our observations also suggest that when multiple EC cells within the same villus or adjacent crypts are activated simultaneously, they collaborate to stimulate the same nerve fibers. Thus, we propose that sparsely distributed EC cells integrate the information within the local environment in a parallel processing manner, converging their signals to produce a singular output to the nervous system inside and outside the gut.

### Mucus as a protective barrier for chemical irritants

The mucus layer is thickest over the crypts, protecting stem cells that regenerate the intestinal epithelium. It therefore makes sense that TRPA1 channels are located preferentially in this protected zone, where they can serve as sentinels for highly reactive irritants such as acrolein, ingestion of which elicits frequent vomiting in dogs (*45*). Acrolein, an environmental toxicant, is found in fried foods and alcoholic beverages and produced by catabolism of cyclophosphamide and related chemotherapeutic agents (*46*). Acrolein may also be produced by microbial metabolism of glycerol in the gut, representing another pathological scenario in which EC cell activation initiates protective nocifensive signals (*46*).

Weaker electrophiles found in foods such as mustard, garlic, and onions do not usually evoke an extreme nocifensive reflex, consistent with our finding that they do not penetrate the mucus layer to stimulate TRPA1 channels on crypt EC cells. In pathological states like colitis or bacterial infection, where the mucus layer is compromised (*47*), these dietary electrophiles could potentially breach this protective barrier to activate crypt EC cells and promote nausea. This is consistent with observations that patients with inflammatory bowel disease (IBD) often avoid spicy foods, including mustard and garlic, suggesting heightened exposure of their crypt EC cells to luminal contents (*48*).

### Detecting stress signals in villi

TRPA1 is activated by reactive oxygen species (ROS) such as H_2_O_2_ and 4-hydroxynonenal, making it a key physiologic sensor of oxidative stress and cellular redox state (*49*). TRPM2 is also activated by H_2_O_2_ and our results suggest that these two TRP channel subtypes function as irritant / ROS sensors in crypts versus villi, respectively (Figure 6F). Unlike TRPA1 receptors in crypts, TRPM2 channels in villi are not as well shielded by a thick mucus layer and may therefore serve as ‘first responders’ to oxidative stress. Moreover, the ability of activated villus EC cells to simultaneously release serotonin and ATP likely augments their capacity to stimulate mucosa-innervating vagal neurons, most or all of which respond to serotonin and ATP. Purinergic receptors are expressed by other cell types in the gut, such as enteric glia (P2X7), enterocytes (P2X7), and secretomotor neurons (P2Y1) (*50*), and it is therefore possible that villus EC cells target these receptors to trigger additional stress responses.

ROS are produced in pathological situations such as IBD and chronic granulomatous disease (*51*). Furthermore, chemotherapeutic drugs can rapidly generate ROS during the initial treatment stages, causing damage to the intestinal mucosa (*51*). Our findings suggest that ROS produced in these patients may activate TRPM2 in villus EC cells, triggering GI pain and nausea. With the loss of the protective mucus layer, TRPA1 channels in the crypt may then be recruited, further contributing to nociceptive and neurogenic inflammatory responses. Several TRPA1 antagonists have undergone clinical trials for managing inflammatory pain or airway hypersensitivity (*52*); our work now highlights their potential use for treating gastrointestinal symptoms associated with overproduction of reactive irritants (of microbial or inflammatory origin) or reduction of the protective mucosal barrier. The same may pertain to potent and selective TRPM2 inhibitors, which are currently lacking.

## Materials and Methods

### Experimental model and subject details Mice

All experimental procedures were conducted in accordance with guidelines approved by the Institutional Animal Care Committees at UCSF, SAHMRI Animal Ethics Committee, and Peking University, and aligned with the NIH Guide for the Care and Use of Laboratory Animals. Subjects were 8-16-week-old mice of both sexes, given ad libitum access to standard lab chow and sterile water. They were housed in a controlled environment under a 12-hour light/dark cycle. For gGRAB_5HT3.0_ sensor imaging, Villin-Cre (Jackson Laboratory. Strain no. 021504) and Insm1-GFP-Cre (gift from Corey Harwell. MMRRC ID: 36986) were crossed to the gGRAB_5HT3.0_-P2A-jRGECO1a reporter line. GCaMP imaging in organoids used Tac1-IRES-Cre (Jackson Laboratory, Strain no. 021877) crossed with GCaMP5g-IRES-tdTomato mice (gift from Lily Jan, Jackson Laboratory, Strain no. 024477). Excitatory DREADD hM3Dq receptors were expressed in EC cells using Tac1-IRES-Cre;ePet1-Flp;FL-hM3Dq mice. Htr3a-GFP mice (MMRRC ID: 000273) visualized 5-HT3 expressing nerve fibers. Pirt1-Cre mice (gift from Xinzhong Dong) were crossed with Ai14 tdTomato reporter mice (Jackson Laboratory, Strain no. 007914) for mucosal dendrite visualization.

### Epithelial cell isolation and organoid culture

Adult male Tac1-IRES-Cre;GCaMP5g-IRES-tdTomato mice were used to generate intestinal organoids as previously reported (*53*), specifically utilizing the upper jejunum to avoid ectopic Tac1-IRES-Cre expression in the lower intestine. Organoids were maintained and passaged every 6 days in organoid growth media (advanced Dulbecco’s modified Eagle’s medium/F12 supplemented with penicillin/streptomycin, 10 mM HEPES, Glutamax, B27 [Thermo Fisher Scientific], 1 mM N-acetylcysteine [Sigma], 50 ng/ml of mouse recombinant epidermal growth factor [Thermo Fisher Scientific], R-spondin1 [10% final volume] and 100 ng/ml of murine Noggin [Peprotech]). For villus organoid differentiation, day 4 organoids were treated with 5 μM IWP2 (Stemgent), 10 μM DAPT (Sigma), 1 μM PD0325901 (Sigma), and 20 ng/mL BMP4 (Peprotech) for 4 days.

### Cell lines

The R-spondin 1 expressing HEK293T (ATCC) cells were maintained in DMEM, 20% fetal calf serum, 1% penicillin/streptomycin, and 125 μg/mL Zeocin (Thermo Fishesr Scientific) at 37°C, 5% CO_2_. Zeocin was removed upon production of R-spondin 1 conditioned media. HEK293T cells (ATCC) were grown in DMEM, 10% fetal calf serum, and 1% penicillin/streptomycin at 37°C, 5% CO_2_ and transfected using Lipofectamine 3000 (Thermo Fisher Scientific) according to manufacturer’s protocol. For biosensor experiments, 200 ng pDisplay-gGRAB_5HT2m_-IRES-mCherryCAAX (Addgene, #208710), 200 ng pcDNA3-5-HT2A-P2A-GCaMP8m, or 200 ng pDisplay-ATP1.0-IRES-mCherryCAAX (Addgene, #167582) was transfected to HEK293T cells in 24-well plates. For 5-HT3 biosensor experiment, 200 ng pcDNA3-5-HT3A and 20 ng pcDNA3-mApple were co-transfected to HEK293T cells in 24-well plates.

## Method details

### Generation of gGRAB_5HT3.0_-P2A-jRGECO1a reporter mouse

The gGRAB_5HT3.0_-P2A-jRGECO1a reporter mouse was generated with help of Biocytogen Pharmaceuticals Co., Ltd. (Beijing, China). In detail, the CAG-loxP-STOP-loxP-gGRAB_5HT3.0_-P2A-jRGECO1a-WPRE-bGH sequence was inserted to the *Rosa26* locus of mouse embryonic stem (ES) cells using CRISPR/Cas9-mediated homology-directed repair (HDR). Successful targeting was confirmed with PCR. The genetically modified ES cells were injected into eight-cell stage embryos to generate chimeric mice. The chimeric mice were then mated with wild-type mice to obtain germline transmission of the targeted allele. The resulting transgenic mouse line stably expresses both the gGRAB_5HT3.0_ sensor and jRGECO1a under the CAG promoter at the *Rosa26* locus upon excision of the floxed stop cassette by Cre recombinase.

### Anti-CD3 antibody-induced inflammation model

Mice received a single intraperitoneal injection of 30 μg anti-CD3 antibody (Thermo Fisher Scientific) diluted to 200 μL with physiological saline. Mice were sacrificed for tissue collection after 12 hours.

### *Ex vivo* serotonin sensor imaging

A ∼1 cm piece of the jejunum was isolated from an 8-16-week old Villin-Cre;gGRAB_5HT3.0_-P2A-jRGECO1a or Insm1-GFP-Cre;gGRAB_5HT3.0_-P2A-jRGECO1a mouse. The isolated tissue was then immediately filleted-open along the mesentery, pinned down to a Sylgard-coated recording chamber, and imaged from the smooth muscle side to observe crypts and from the luminal side to observe villi. Imaging was performed with a Leica SP8 confocal microscope with an HC APO L 20x/1,00 W objective and LAS X software (Leica Microsystems). The tissue was bath perfused with bubbled room-temperature Krebs buffer (118 mM NaCl, 4.7 mM KCl, 1 mM MgCl_2_, 2 mM CaCl_2_, 1.2 mM KH_2_PO_4_, 25 mM NaHCO_3_, and 10 mM D-glucose) at a rate of ∼1 mL/min. All pharmacological reagents were diluted in Krebs buffer and bath perfused with simultaneous manual application. For recordings of StcE digested tissues, 50 μM StcE was supplied to Krebs buffer. At the end of each recording, the gGRAB_5HT3.0_ sensor was fully activated by bath applied 20 μM serotonin. Acquired images were analyzed with Fiji software v2.14 (NIH). The regions of interest (ROIs) were drawn around individual crypts or villi and ΔF/F_0_ was calculated and normalized to serotonin-activated maximum signals. The area under the curve (A.U.C.) was calculated as (Normalized gGRAB_5HT3.0_ΔF/F_0_) for the duration of 5 minutes during baseline or drug application. When measuring the baseline serotonin levels in villi, 20 μM RS 23597-190 was added at the end of recordings to fully quench the sensor.

### Serotonin sensor imaging with isolated villi and crypts

Pieces of the jejunum were isolated from an 8-16-week-old Villin-Cre;gGRAB_5HT3.0_-P2A-jRGECO1a mice. The isolated tissue was then filleted-open along the mesentery. For recordings from isolated villi, villi were scraped off using glass coverslips and resuspended in a 50% Matrigel-Krebs buffer mixture. Matrigel domes (∼5 μL) were then formed on glass coverslips for imaging. The villi exposed to the surface of the Matrigel domes were identified under a microscope and used for imaging. For recordings from isolated crypts, after villi removal, the tissue was incubated in 10 mL of cold Dulbecco’s Phosphate-Buffered Saline (DPBS) with 30 mM EDTA for 20 minutes, followed by vigorous shaking for 30-60 seconds. Isolated crypts were filtered through 70 μm strainers and plated onto CellTak (Corning)-coated coverslips. Serotonin sensor imaging was performed with an upright microscope equipped with a Grasshopper 3 (FLIR) camera and a Lambda LS light source (Sutter). Villi and crypts were maintained under a constant laminar flow of Ringer’s solution applied by a pressure-driven microperfusion system (SmartSquirt, Automate Scientific). All pharmacological reagents were delivered by local perfusion. Acquired images were analyzed with Fiji software (NIH). ROIs were drawn around individual EC cell and ΔF/F_0_ was calculated.

### Expression and purification of StcE

The pET28b-StcE-Δ35-NHis plasmid (gift from Carolyn Bertozzi) was transformed into *E*.*coli* BL21(DE3). Transformed *E*.*coli* cells were cultured in LB medium containing 50 μg/L kanamycin at 37°C for 4 hours. Isopropyl-thio-β-D-galactopyranoside (IPTG) was added to a final concentration of 0.3 mM to induce protein expression. Following an additional incubation at 20°C for 12 hours, cells were harvested by centrifugation and resuspended in purification buffer (500 mM NaCl and 20 mM HEPES-Na [pH 7.5]). Cell extracts were obtained by sonication followed by centrifugation at 36,000 g for 30 minutes. The supernatant was incubated with 2 mL Ni-NTA (Qiagen) for 1 hour at 4°C with gentle mixing. The resin was washed in batch with 5 column volumes of purification buffer, then loaded onto a column and further washed with 5 column volumes of purification buffer + 20 mM imidazole and 10 column volumes of purification buffer + 30 mM imidazole. The column was then eluted with purification buffer + 250 mM imidazole. To remove imidazole, the eluted protein was concentrated to 30 mM and then dialyzed against purification buffer overnight. Purified proteins are then stored at 4°C.

### Mucus digestion with StcE

Purified StcE (30 mM) was diluted to 10 mM with double distilled H_2_O and 1 M HEPES-Na (pH 7.5) solution was added to a final concentration of 20 mM. This dilution was performed immediately before the experiment to avoid precipitation of StcE. The isolated jejunum (∼1 cm) was incubated in 10 mL of 10 mM StcE for 60 minutes at room temperature with gentle shaking. The StcE solution was exchanged after 30 minutes. Digested tissues were immediately mounted on a recording chamber for gGRAB_5HT3.0_ sensor imaging.

### BODIPY-IA staining

A 10 mM StcE solution was prepared as described above. Sections of the jejunum (∼1 cm) were moved to 10 mL of a solution of 166 mM NaCl + 20 mM HEPES-Na (pH 7.5) with or without 10 mM StcE, and incubated for 30 minutes at room temperature with gentle shaking. Digested tissues were immediately transferred to 10 μM BODIPY-FL-iodoacetamide (Thermo Fisher Scientific) in Dulbecco’s Phosphate-Buffered Saline (DPBS) and incubated for 3-15 minutes at room temperature with gentle shaking. Stained tissues were briefly rinsed with DPBS and fixed with 4% paraformaldehyde (PFA) for 3 hours at 4°C. Fixed tissues were dehydrated in 30% sucrose overnight at 4°C. The tissues were embedded in Tissue-Tek O.C.T. Compound (Sakura Finetek USA) and subsequently sectioned at a thickness of 10 μM on a Leica CM3050 S cryostat. The nuclei were stained with 4,6-diamidino-2-phenylindole (DAPI, 0.5 μg/ml, Thermo Fisher Scientific) and sections were mounted with ProLong Diamond Antifade Mountant (Thermo Fisher Scientific). Confocal images were captured on an inverted Nikon Ti microscope run using Micro Manager 2.0 Gamma (*54*), equipped with a Zyla 4.2 CMOS camera (Andor), piezo XYZ stage (ASI), CSU-W1 Spinning Disk with Borealis upgrade (Yokogowa/Andor), Spectra-X (Lumencor), ILE 4 line Laser Launch (405/488/561/640 nm; Andor). Images were taken using a Plan Apo λ 20x / 0.75 using lasers 405, 488, and 561 nm and emission filters 447/60, 525/50, 607/36, for DAPI, GFP, and RFP, respectively. Maximum intensity projections were generated in Fiji v2.14.

### Expression and purification of the 5-HT3 nanobody

The 5-HT3 nanobody (VHH15, gift from Hugues Nury) was engineered to include a C-terminal fusion with mCherry-His6 through an SSGSS linker and a gp64 signal peptide sequence was appended to the N-terminus, facilitating secretion into the insect cell culture media. The resultant plasmid, pFastBac-gp64-VHH15-mCherry-His6, was used to transfect Sf9 cells, generating P1 virus. Sf9 cells were then infected with amplified P2 virus for nanobody expression and harvested at 60 hours post-infection. The culture was centrifuged at 3,500 g for 15 minutes and the supernatant was filtered through a 0.22 μm filter. The filtered supernatant was adjusted to pH 7.5 using 1 M HEPES (pH 8.0), and divalent ions were replenished by adding 5 mM CaCl_2_ and 2 mM NiSO_4_. The supernatant was then incubated with 2 mL of Ni-NTA resin for 2 hours at 4°C. The resin was washed in batch with 5 column volumes of VHH15 buffer (500 mM NaCl and 50 mM Tris [pH 8.0]), then loaded onto a column and further washed with 5 column volumes of VHH15 buffer with 20 mM imidazole. The column was then eluted with 3 column volumes of elution buffer (125 mM NaCl, 250 mM imidazole, and 50 mM Tris [pH 7.4]). The elute was concentrated and loaded onto a Superdex 200 10/30 (GE Healthcare) gel filtration column in the buffer 10 mM HEPES (pH 7.5) and 100 mM NaCl. Fractions containing the peak were pooled and concentrated to 10 μM (0.39 mg/mL).

### Histology and immunostaining

Immunofluorescence in the small and large intestine was performed using 10 μm cryosections. Blocking was performed with 5% w/v BSA (Sigma), 5% normal serum corresponding to secondary antibody species, and 0.3% Triton-X in PBS at room temperature for 30 minutes. Primary antibodies were incubated overnight at 4°C at the indicated dilutions. Antibodies used were against serotonin (1:5,000, Immunostar), Tuj1 (1:500, Abcam), Collagen IV (1:500 Abcam), GFP (1:500, Abcam), and mCherry (1:500, Takara). Secondary antibodies from Invitrogen (Alexa Fluor 647 goat anti-rabbit, Alexa Fluor 568 goat anti-rabbit, and Alexa Fluor 488 goat anti-chicken) were incubated at 1:500 dilution for 2 hours at room temperature at 1:500 dilution. Z-stack images were taken with a Nikon CSU-W1 spinning disk confocal microscope as described above (UCSF Center for Advanced Light Microscopy). Maximum intensity projections were generated in Fiji v2.14.

### Whole-mount staining of EC cells and mucosal dendrites

A piece of the jejunum was isolated from 8-16 week-old Pirt1-Cre;Ai14 mice. Pirt1-Cre;Ai14 mice were used because of the high expression level of tdTomato in mucosal dendrites. Isolated tissues were washed, filleted-open, and fixed with 4% PFA for 4 hours at 4°C. Blocking was performed with 5% w/v BSA (Sigma), 5% donkey serum, and 0.3% Triton-X in PBS at room temperature for 3 hours. Tissues were then incubated in a primary antibody solution (1:500 rabbit anti-serotonin, Immunostar, and 1:500 goat anti-collagen IV, abcam) for two days at 4°C. Following primary incubation, tissues were washed three times in PBS with 0.2% Triton-X and incubated overnight in a secondary antibody solution (1:500 Alexa Fluor 488 donkey anti-rabbit and Alexa Fluor 647 donkey anti-goat). After overnight, tissues were washed three times in PBS with 0.2% Triton-X and mounted with ProLong Diamond Antifade Mountant. Z-stack images were taken with a Nikon CSU-W1 spinning disk confocal microscope as described above, using a Plan Apo VC 100x / 1.4 Oil objective (UCSF Center for Advanced Light Microscopy). Image deconvolution and 3D image reconstruction were conducted using Huygens (Scientific Volume Imaging) and Imaris (Oxford Instruments), respectively. The distances between the basolateral side of EC cells and the nearest dendrites were manually measured in Imaris.

### Immunostaining of HEK293T cells with the 5-HT3 nanobody

HEK293T cells were plated on 4-well chamber slides (ibidi) and transfected with either pcDNA3-5-HT3A or the empty pcDNA3 plasmid. After overnight, cells were fixed with 4% PFA for 20 minutes at room temperature. Fixed cells were incubated with VHH15-mCherry in PBS + 0.1% Triton-X (1:20 dilution) for 1 hour at room temperature. After staining, cells were washed with PBS + 0.1% Triton-X three times and mounted with ProLong Diamond Antifade Mountant. Z-stack images were taken with a Nikon CSU-W1 spinning disk confocal microscope as described above (UCSF Center for Advanced Light Microscopy). Maximum intensity projections were generated in Fiji v2.14.

### Dorsal root ganglia (DRG) isolation and immunostaining with the 5-HT3 nanobody

DRGs were harvested from both male and female Htr3a-EGFP mice between 8-16 weeks of age. The dissected DRGs were fixed in 4% PFA for 3 hours at 4°C. Blocking was performed with 5% w/v BSA (Sigma), 5% goat serum, and 0.3% Triton-X in PBS at room temperature for 3 hours. The DRGs were then incubated with VHH15-mCherry (1:20 dilution) in PBS with 0.1% Triton-X overnight at 4°C. Post incubation, the DRGs were briefly washed in PBS and fixed again in 4% PFA for 30 minutes at 4°C. Following a PBS wash, DRGs were transferred to a primary antibody solution (1:500 chicken anti-GFP, abcam, and 1:500 rabbit anti-mCherry, Takara) overnight. DRGs were then washed three times in PBS with 0.2% Triton-X, and moved to a secondary antibody solution overnight (1:500 Alexa Fluor 488 goat anti-chicken and Alexa Fluor 568 goat anti-rabbit). The stained DRGs were then washed three times in PBS with 0.2% Triton-X and mounted with ProLong Diamond Antifade Mountant. Z-stack images were taken with a Nikon CSU-W1 spinning disk confocal microscope as described above (UCSF Center for Advanced Light Microscopy). Maximum intensity projections were generated in Fiji v2.14.

### Immunostaining of the intestine with the 5-HT3 nanobody

The jejunum and proximal colon were harvested from male and female *Htr3a*-EGFP mice between 8-16 weeks of age. Tissues were filleted-open, washed, and incubated in the staining buffer containing VHH15-mCherry (1:30 dilution) and protease inhibitor cocktail (Roche) in Krebs buffer for 2 hours at room temperature. After staining, the tissues were washed three times with DPBS and fixed in 4% PFA for 1 hour at room temperature. Blocking was performed with 5% w/v BSA (Sigma), 5% goat serum, and 0.3% Triton-X in PBS at room temperature for 3 hours. Tissues were then incubated in a primary antibody solution (1:500 chicken anti-GFP, abcam, and 1:500 rabbit anti-mCherry, Takara) for two days at 4°C. Following incubation, the tissues were washed three times in PBS with 0.2% Triton-X, and incubated in a secondary antibody solution (1:500 Alexa Fluor 488 goat anti-chicken and Alexa Fluor 568 goat anti-rabbit). After an overnight incubation, the tissues were washed three times in PBS with 0.2% Triton-X and mounted with ProLong Diamond Antifade Mountant. Z-stack images were taken with a Nikon CSU-W1 spinning disk confocal microscope as described above (UCSF Center for Advanced Light Microscopy). Maximum intensity projections were generated in Fiji v2.14.

### *In situ* hybridization

10 μm cryosections were prepared as described above. Single-molecule RNA-FISH was performed using the RNAscope Multiplex Fluorescent Detection Kit v2 (Advanced Cell Diagnostics) according to manufacturer’s protocol. Z-stack images were taken with a Nikon CSU-W1 spinning disk confocal microscope as described above (UCSF Center for Advanced Light Microscopy). Maximum intensity projections were generated in Fiji v2.14.

### GCaMP imaging using intestinal organoids

Five days after passage, Tac1-IRES-Cre;GCaMP5g-IRES-tdTomato organoids were removed from Matrigel (Corning) and mechanically broken up with a 1000 μL pipette. The organoid fragments were seeded onto Cell-Tak (Corning)-coated coverslips and placed in a recording chamber containing Ringer’s solution (140 mM NaCl, 5 mM KCl, 2 mM CaCl_2_, 2 mM MgCl_2_, 10 mM D-glucose, and 10 mM HEPES-Na [pH 7.4]). EC cells were identified by tdTomato expression. GCaMP imaging was performed with an upright microscope equipped with a Grasshopper 3 (FLIR) camera and a Lambda LS light source (Sutter). Organoids were maintained under a constant laminar flow of Ringer’s solution applied by a pressure-driven microperfusion system (SmartSquirt, Automate Scientific). All pharmacological reagents were delivered by local perfusion. Acquired images were analyzed with Fiji software (NIH). ROIs were drawn around individual EC cell and ΔF/F_0_ was calculated.

### Organoid swelling assay

Organoids were first passaged into organoid culture media lacking N-acetylcysteine to prevent the inhibition of TRPA1 channels. 24 hours post-passage, organoids were treated with either 1 μM serotonin, 10 μM RS 23597-190, or 5 μM A967079 in N-acetylcysteine-free organoid culture media. Images of the organoids were captured at 60 minutes post-serotonin treatment and 12 hours post-A967079 treatment. Cross-sectional areas of the organoids were subsequently measured using Fiji v2.14.

### Biosensor experiments

HEK293T cells transiently transfected with biosensor plasmids (gGRAB_5HT2m_ sensor, 5-HT2A receptor and GCaMP8m, 5-HT3 channel, or gGRAB_ATP1.0_ sensor) were dissociated with trypsin and washed once with Ringer’s. The dissociated cells were plated on top of intestinal organoids. Individual HEK293T cells were carefully lifted from coverslips and positioned 5 μm from an EC cell using a glass pipette. For the 5-HT3 biosensor experiments, whole-cell configuration was achieved before lifting the cell. The membrane potential was held at -80 mV to measure inward 5-HT3 currents. For 5-HT2A and gGRAB_ATP1.0_ biosensor experiments, imaging was performed using an upright microscope equipped with a Grasshopper 3 camera (FLIR) and a Lambda LS light source (Sutter Instrument). The entire area of each biosensor cell was used for the calculation of ΔF/F_0_ values. For gGRAB_5HT2m_ biosensor experiments, imaging was performed on a Leica SP8 confocal microscope with LAS X software (Leica Microsystems). At the end of each recording, the gGRAB_5HT2m_ sensor was fully activated with 500 μM serotonin. In these experiments, only the portion of each biosensor cell membrane within 5 μm of an EC cell was used for the calculation of ΔF/F_0_, which was then normalized to the maximum signals activated by serotonin. In all the biosensor experiments, the bath solution was static to prevent the washout of endogenously released serotonin, and pharmacological agents were applied manually with a 1000 μL pipette. All images were analyzed using Fiji v2.14.

### Electrophysiology

Electrophysiological recordings were performed with an Axopatch 200B amplifier (Molecular Devices) connected to Digidata 1550B (Molecular Devices), sampling at 10 kHz and filtering at 1 kHz. Membrane potentials were corrected for liquid junction potentials. Patch electrodes (3-6 MΩ) were pulled from borosilicate capillaries (BF-150-110-10, Sutter Instrument). The external solution for both EC cell and 5-HT3 channel recordings was Ringer’s solution. For TRPM2 recordings, the external solution contained 150 mM NaCl, 2 mM CaCl_2_, 1 mM MgCl_2_, 5 mM HEPES-Na (pH 7.4), and 10 mM D-glucose. The NMDG external solution was composed of 150 mM NMDG-Cl, 2 mM CaCl_2_, 1 mM MgCl_2_, 5 mM HEPES-Na (pH 7.4), and 10 mM D-glucose. The intracellular solution for EC cell recordings consisted of 140 mM K-aspartate, 13.5 mM NaCl, 1.6 mM MgCl_2_, 0.09 mM EGTA, 9 mM HEPS-K (pH 7.35), 14 mM phosphocreatine-tris, 4 mM MgATP, 0.3 mM Na_2_GTP. Intracellular solution for 5-HT3 recordings consisted of 140 mM K-gluconate, 5 mM NaCl, 1 mM MgCl_2_, 10 mM EGTA-K, and 10 mM HEPES-K (pH 7.2). For TRPM2 recordings, the intracellular solution included 150 mM NaCl, 5 mM HEPES-Na (pH 7.4), 5 mM EGTA-Na, 1 mM MgCl_2_, 5.1 mM CaCl_2_ (yielding a final free Ca^2+^ concentration of 100 μM), and 500 μM ADP-ribose (Sigma).

### EC cell dissociation

Enterochromaffin (EC) cells were isolated from the upper half of the small intestine of 8-16-week-old Tac1-IRES-Cre;GCaMP5g-IRES-tdTomato mice. The tissue was cut into approximately 3 cm segments, incubated in 10 mL of cold DPBS with 30 mM EDTA and 1.5 mM DTT on ice for 20 minutes, then transferred to 6 mL of warm DPBS with 30 mM EDTA and incubated at 37°C for 8 minutes. To dissociate the epithelial layer, vigorous shaking was applied for 30-60 seconds. The dissociated epithelium was centrifuged and washed with DPBS containing 10% fetal bovine serum (FBS). The washed epithelium was digested in 10 mL of digestion buffer (HBSS with 0.3 mg/mL dispase II [Sigma] and 0.2 mg/mL DNaseI [Sigma]) at 37°C for 8 minutes, with vigorous shaking at 2-minute intervals. The cells were then washed once with HBSS containing 10% FBS and 0.2 mg/mL DNaseI, filtered through 70 μm and 40 μm strainers, and resuspended in DMEM supplemented with 10% FBS, B27, and 5 μM Y-27632 (Sigma). Cells were plated onto glass coverslips precoated with 5% Matrigel solution. Two days following dissociation, EC cells exhibiting the characteristic polygonal or cone-shaped morphology were predominantly surviving cells from the crypt regions. These crypt-originating EC cells were subsequently utilized for electrophysiological recordings and GCaMP imaging conducted 2-3 days post-dissociation.

### Intestinal Isc measurement in mice

The ileum was excised under anesthesia and soaked in isoosmolar solution containing 300 mM mannitol and 10 μM indomethacin. Mucosa was stripped from serosa/muscle layers under a dissection microscope and mounted on Ussing chambers containing Ringer’s solution (120 mM NaCl, 5 mM KCl, 1 mM CaCl_2_, 1 mM MgCl_2_, 10 mM D-glucose, 5 mM HEPES-Na [pH 7.4], and 25 mM NaHCO_3_) on the basolateral side. For apical side, a similar solution was used except 120 mM NaCl was replaced with 60 mM NaCl and 60 mM sodium gluconate, and glucose was replaced with 10 mM mannitol. Compounds were added to both apical and basolateral bathing solutions unless specified otherwise. The solutions were aerated with 95% O_2_ /5% CO_2_ and maintained at 37°C during experiments. Short-circuit current (Isc) was measured using an EVC4000 multichannel voltage clamp (World Precision Instruments, Sarasota, FL, USA) via Ag/AgCl electrodes and 3 M KCl agar bridges as previously described (*55*).

### Retrograde tracing of mucosal afferents from the proximal small intestine

Adult male mice of C57BL/6 background (Jackson Laboratory) aged 16-20 weeks were used. Retrograde tracing using cholera toxin subunit B (CTB, 0.5%) directly conjugated to Alexa Fluor 488 (Invitrogen, Thermo Fisher Scientific, #C2284, Australia) was performed from the lumen of the proximal small intestine (jejunum) using a technique modified from Harrington et al (*39*). A small aseptic abdominal incision was made in mice anesthetized with isoflurane (2-4% in oxygen). The proximal small intestine was located, and injections of 5 μL were made through the intestinal wall into the lumen at three sites covering a length of 5 cm. The tracer was expelled completely prior to the needle withdrawal back through the intestine wall, which was gently rubbed together using cotton tip applicators to distribute the tracer throughout the lumen. Injections were made with a 30-gauge needle (HAMC7803-07, point style: 4; Hamilton Company, Bio-Strategy, Campbellfield, Vic, Australia) attached to Hamilton 5 μL syringe (HAMC7634-01, 5 μL 700 series RN syringe; Hamilton Company, Bio-Strategy). The abdominal incision was then sutured closed, analgesic (buprenorphine, 0.1 mg/kg) and antibiotic (ampicillin, 50 mg/kg) administration were given subcutaneously as mice regained consciousness. Mice were then housed individually and closely monitored for 4 days prior to nodose collection. Imaging studies revealed the vast majority of these mucosal traced neurons were located within the nodose ganglia rather than the dorsal root ganglia.

### Calcium imaging of dissociated nodose ganglia neurons

Retrogradely traced nodose ganglion neurons were isolated from adult mice. Briefly, 4 days after retrograde tracing, mice were euthanized by CO2 inhalation and nodose ganglia were surgically removed and were digested with 4 mg/mL collagenase II (GIBCO, Life Technologies) plus 4 mg/mL dispase (GIBCO) for 30 min at 37°C, followed by 4 mg/mL collagenase II for 10 min at 37°C, similar to that described previously for DRG (*6*). Neurons were then mechanically dissociated into a single-cell suspension via trituration through fire-polished Pasteur pipettes. Neurons were resuspended in DMEM (GIBCO) containing 10% FCS (Invitrogen), 2mM L-glutamine (GIBCO), 100 mM MEM non-essential amino acids (GIBCO), 100 mg/ml penicillin/streptomycin (Invitrogen) and 100 ng/ml NGF (Sigma). Neurons were spot-plated on coverslips coated with poly-D-lysine (800 mg/ml) and laminin (20 mg/ml) and maintained at 37°C in 5% CO_2_. After 24 h in culture, neurons were loaded with 2.5 μM Fura-2-AM (Thermo Fisher Scientific) and 0.02% (v/v) pluronic acid for 30 min at room temperature in Ringer’s solution ((NaCl 140 mM, KCl 5 mM, CaCl2 1.25 mM, MgCl2 1 mM, glucose 10 mM, HEPES 10 mM, pH 7.4). After a brief wash, coverslips were transferred to a recording chamber filled with Ringer’s solution at room temperature (about 22°C). Retrogradely traced nodose ganglion neurons were identified by the presence of the 488 tracer and viability was verified by responses to 40 mM KCl. Fura-2-AM fluorescence was measured at 340 nm and 380 nm excitation, and 530 nm emission was measured using an Olympus IX71 microscope in conjunction with a Sutter Lambda 10-3 wavelength switcher and the Chroma filter set no. 49011 (ET480/40x (Ex), T510lpxrxt (BS), ET535/50m (Em)). Fluorescence images were obtained every 5 seconds, using a ×4 objective with a monochrome CCD camera (Retiga ELECTRO). Images were taken at baseline and following administration of Adenosine 5’-triphosphate disodium salt (ATP, 10 μM, Sigma Merck), m-Chlorophenylbiguanide hydrochloride (mCPBG, 10 μM, Tocris), Tegaserod maleate (1 μM, Sigma Merck), Capsaicin (50 nM, Sigma Merck), Allyl isothiocyanate (AITC, 1 μM, Sigma Merck), H_2_O_2_ (.00 μM, Sigma Merck) and KCl (40 mM). Fluorescence traces of cell bodies were extracted using Metafluor software (Molecular Devices). Regions of Interest (ROIs) were manually drawn around the cell bodies of neurons and their fluorescence traces were extracted as 340/380 ratio.

### Single-cell RT-PCR on mucosal traced nodose ganglion neurons

Four days after the mucosal retrograde tracing procedure described above, nodose ganglia were removed and enzymatically dissociated. Neurons were allowed to settle for two hours before adding 2 mL of media and preparing the coverslips for single-cell picking. Traced cells were manually picked using a micromanipulator under a microscope equipped with an appropriate fluorescent filter. Cells were under a continuous slow flow of RNA/DNAse free PBS to reduce potential contamination. After picking a traced cell, the glass capillary was broken into a tube containing 9 μL lysis buffer with 1 μL DNAse I (TaqMan^®^ Gene Expression Cells-to-CT™ Kit; Thermo Fisher Scientific). A bath control was taken and analyzed from every coverslip with other samples. The whole cell lysate was used for cDNA synthesis using SuperScript™ VILO™ Master Mix (Thermo Fisher Scientific) and diluted 1:5 for further PCR analysis. PCR was performed according to the manufacturer’s instructions using TaqMan™ Gene Expression Master Mix (Thermo Fisher Scientific) for 55 cycles. A target was defined to be present when a typical amplification curve was produced. Predesigned Taqman probes were purchased from Thermo Fisher Scientific (*P2rx2*, Mm00462952_m1; *P2rx3*, Mm00523699_m1; *Htr3a*, Mm00442874_m1; *Htr3b*, Mm00517424_m1; *Htr4*, Mm00434129_m1; *Trpv1*, Mm01246300_m1; *Trpa1* Mm01227437_m1 and *Trpm2*, Mm00663098_m1).

## Statistical analysis

Data were analyzed with Prism (Graphpad) and n represents the number of cells, crypts, villi, or independent experiments. Data were considered significant if p < 0.05 using paired or unpaired two-tailed Welch’s t-test or one-way ANOVAs. Statistical parameters are described in figure legends. All significance tests were justified considering the experimental design and we assumed normal distribution and variance, as is common for similar experiments. Sample sizes were chosen based on the number of independent experiments required for statistical significance and technical feasibility.

## Data availability

All data generated or analyzed during this study are included in the manuscript and supplementary information. Reagents and codes used here are available upon request.

## Acknowledgements

We thank Ms. Jeannie Poblete for her technical support. We thank Drs. James Bayrer and Holly Ingraham, and all members of the Julius lab for discussion. We thank Dr. Hugues Nury for sharing the VHH15 plasmid. We thank Dr. Carolyn Bertozzi for sharing the StcE plasmid. We appreciate the support from staff in UCSF’s core facilities, including the Center for Advanced Light Microscopy (Drs. DeLaine Larsen, Kari Herrington, and So Yeon Kim, S10 Shared Instrumentation Grant 1S10OD017993-01A1 for Nikon CSU-W1 spinning disk confocal microscope). This work was supported by NIH grants NS105038, NS113869, and DK135714 to D.J.; BRAIN Initiative 1U01NS113358 and 1U01NS120824 to Y.L.; DK126070, DK072517, and EY036139 to O.C., the National Natural Science Foundation of China (31925017 to Y.L.), the New Cornerstone Science Foundation through the New Cornerstone Investigator Program and the XPLORER PRIZE to Y.L., National Health and Medical Research Council (NHMRC) of Australia Investigator Leadership Grant (APP2008727) to S.M.B., and Cystic Fibrosis Foundation to O.C.. K.K.T. was supported by a Damon Runyon Cancer Research Foundation Fellowship (DRG-[2387-30]).

## Author contributions

Research design: K.K.T., O.C., S.M.B., Y.L., and D.J. Writing: K.K.T. and D.J., with input from all authors. Patch-clamp recordings of EC cells: K.K.T. and N.D.R. Development of gGRAB_5HT3.0_: F.D. Ussing Chamber: T.C. Mucosal retrograde tracing: A.M.H. Real-time PCR and analysis: S.G.C. Ca^2+^ imaging and analysis of vagal neurons: M.B. and T.O. All other experiments and analysis: K.K.T. Supervision of trainees: O.C., S.M.B., Y.L., and D.J. Project administration: K.K.T., O.C., S.M.B., Y.L., and D.J.

---

**Supplementary Figure 1.**
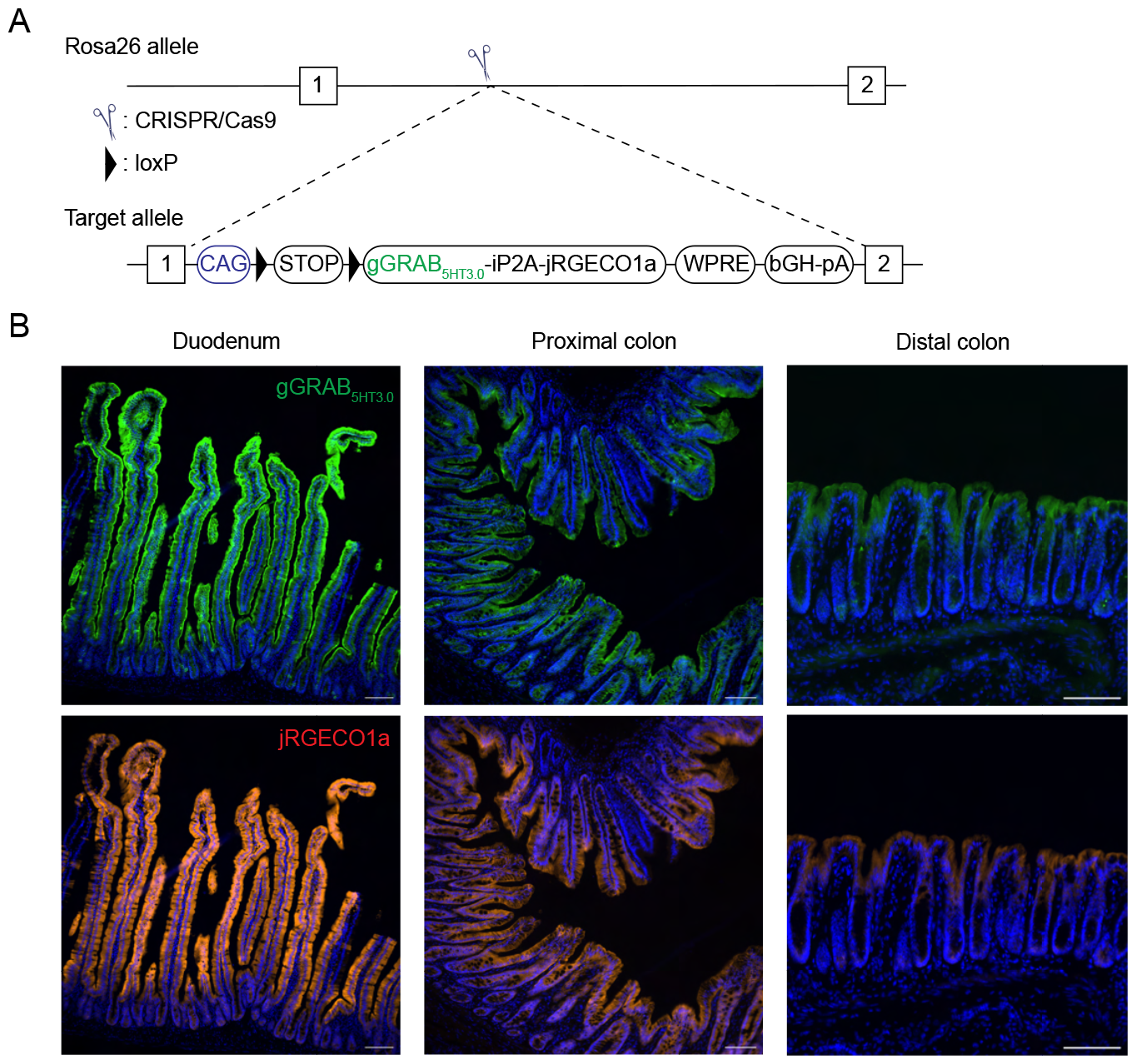
(A) Strategy to generate the transgenic knock-in mouse line expressing gGRAB_5HT3.0_ and jRGECO1a in the *Rosa26* locus. (B) Immunohistochemical labeling of gGRAB_5HT3.0_ sensors and jRGECO1a in Vil1-Cre;gGRAB_5HT3.0_ mice. gGRAB_5HT3.0_ and jRGECO1a were immunostained with an anti-GFP and anti-mCherry antibody, respectively. Scale bar = 100 μm.

**Supplementary Figure 2.**
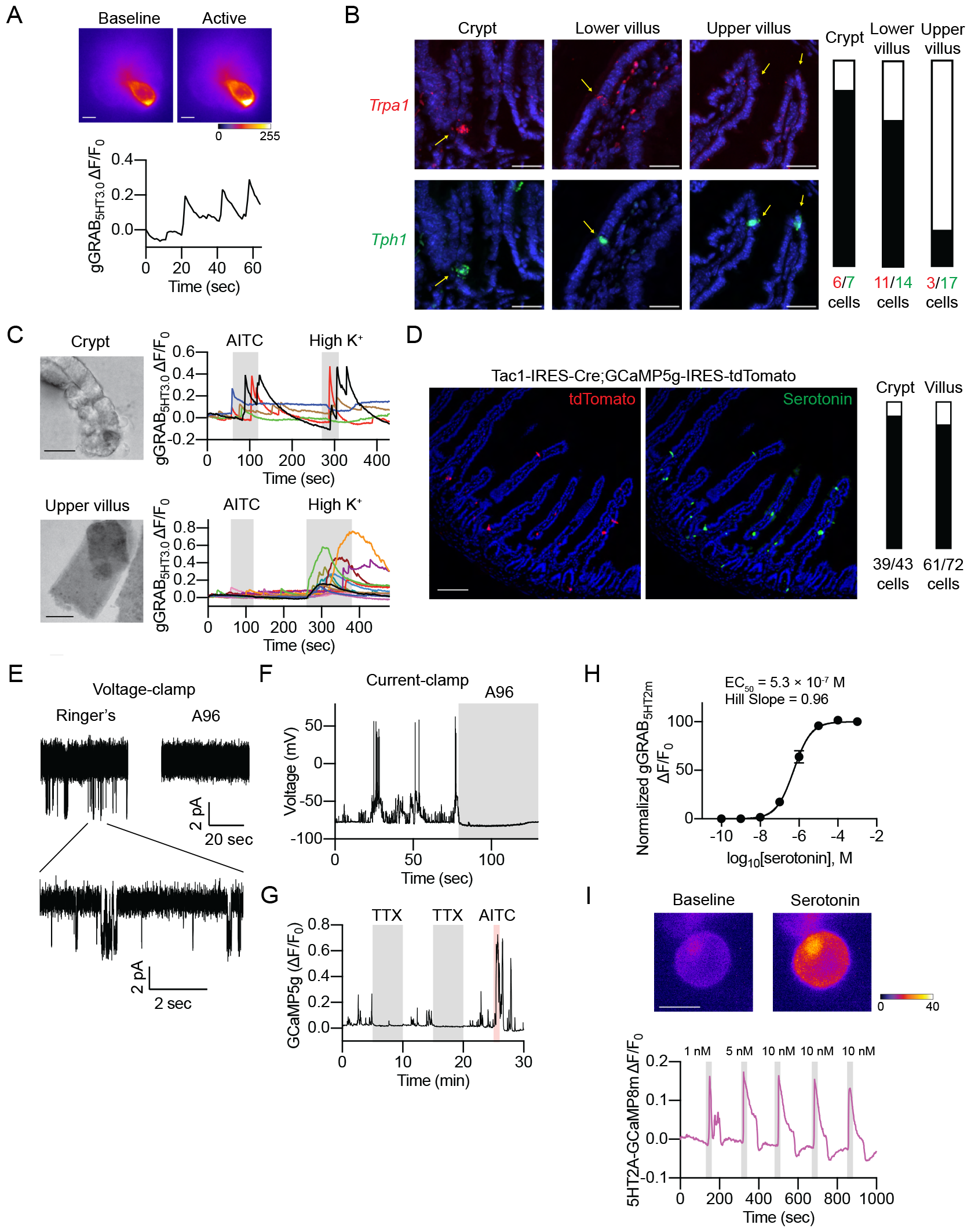
(A) Spontaneous serotonin release in dissociated crypts from the gGRAB_5HT3.0_ sensor mice. Scale bar = 5 μm. (B) *In situ* hybridization of *Trpa1* and *Tph1* in the small intestine. *Tph1* is the rate-limiting enzyme in the serotonin synthesis that is expressed in EC cells in both crypts and villi. *Trpa1* expression is predominantly limited to the crypts and lower villi. Bars indicate the ratio of *Trpa1*-positive cells / *Tph1*-positive cells. Scale bar = 30 μm. (C) AITC activates crypt but not upper villus EC cells in primary isolated crypts and villi from gGRAB_5HT3.0_ sensor mice. Scale bar = 30 μm. (D) Immunohistochemical labeling of crypt and villus EC cells in Tac1-IRES-Cre;GCaMP5g-IRES-tdTomato mice. Bars indicate the ratio of tdTomato-positive cells / serotonin-positive cells. Scale bar = 100 μm. (E) (Left) A96-sensitive spontaneous TRPA1 channel opening in a crypt EC cell. The membrane potential was held at -80 mV. (F) In the same cell, A96 inhibits the spontaneous action potentials in the same EC cell. 10 μM A96 was applied as indicated above the signal. (G) Tetrodotoxin (TTX) inhibits the spontaneous GCaMP activity of EC cells. (H) Normalized dose-response curve of gGRAB_5HT2m_ for serotonin in the presence of 1 μM citalopram, a serotonin transporter inhibitor (EC_50_ = 5.3 × 10^-7^ M. Hill Slope = 0.96. n = 10). (I) GCaMP imaging of HEK cells expressing 5-HT2A receptors and GCaMP8m. The biosensor cells repeatedly responded to 1-10 nM serotonin. Scale bar = 10 μm.

**Supplementary Figure 3.**
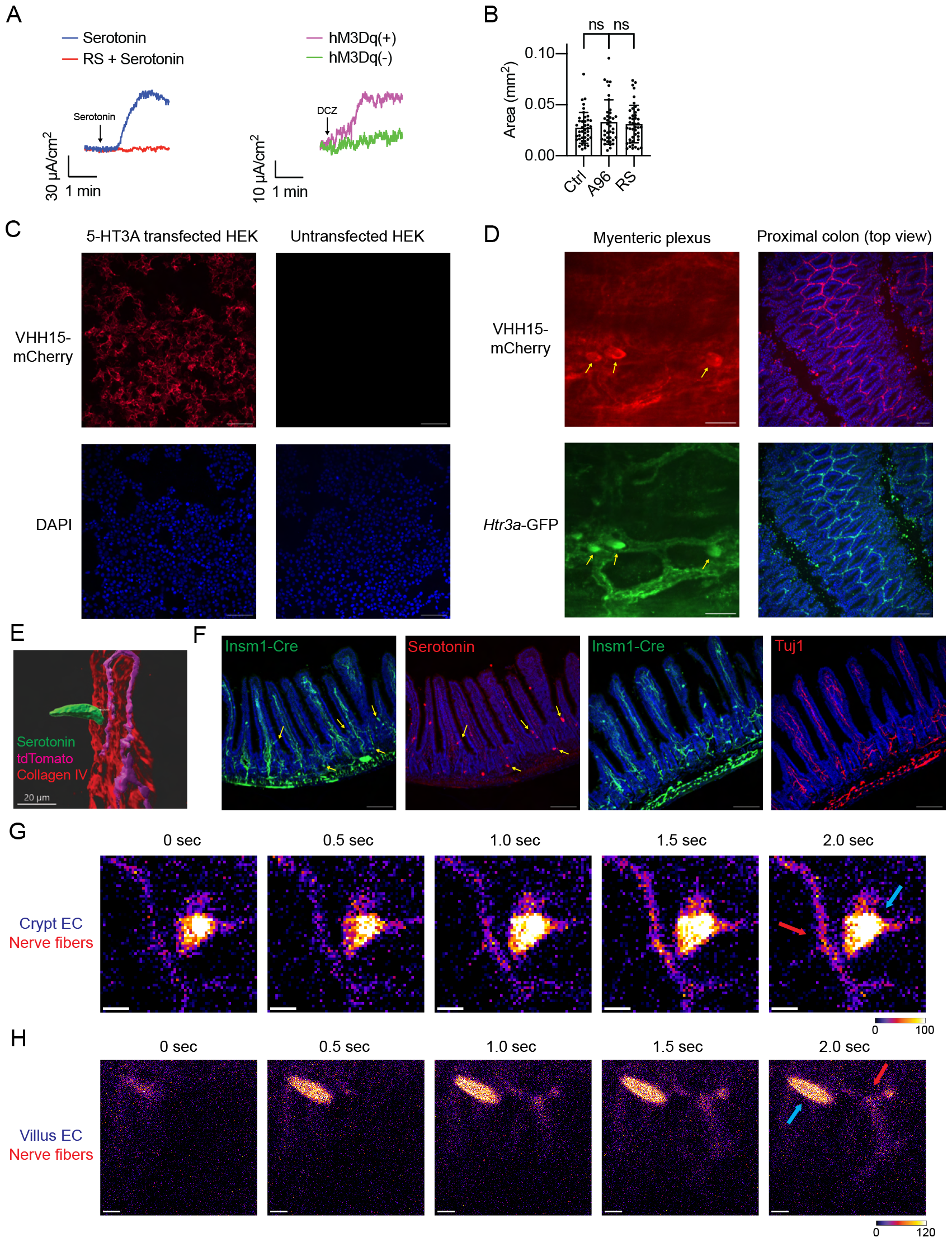
(A) (Left) Time course of short-circuit currents (Isc) in response to 1 μM serotonin with or without a 5-HT4 antagonist (1 μM RS 23597-190). (Right) Time course of short-circuit currents (Isc) in response to a DREADD receptor agonist, 1 μM DCZ, in the ileal mucosa isolated from Tac1-Cre;ePet-Flp;hM3Dq (hM3Dq(+)) or hM3Dq(-) control mice. (B) Intestinal organoids were grown in the presence of 5 μM A96 or 10 μM RS for 4 days, and cross-sectional areas were compared between untreated organoids. n = 38-44. Ordinary one-way ANOVA; ns = not significant. (C) Immunohistochemical labeling of HEK cells transiently transfected with 5-HT3A. Transfected or untransfected HEK cells were stained with VHH15-mCherry. Scale bar = 100 μm. (D) Immunohistochemical labeling of 5-HT3 receptors in the myenteric plexus and proximal colon with VHH15-mCherry. The signal was enhanced by staining for mCherry. Yellow arrows indicate cell bodies of *Htr3a*-GFP(+) neurons in the myenteric plexus. Scale bar = 50 μm. (E) Three-dimensional rendering of EC cell (green), nerve fibers (purple), and basolateral membrane (red). Scale bar = 20 μm. (F) Insm1-Cre labels both EC cells and mucosal nerves. EC cells were labeled with an anti-serotonin antibody and mucosal nerves were labeled with an anti-Tuj1 antibody. Yellow arrows indicate EC cells. Scale bar = 100 μm. (G) and (H) Time-lapse images depicting the diffusion of serotonin, derived from (G) crypt EC cells and (H) villus EC cells, onto the mucosal nerve fibers. Blue and red arrows indicate EC cells and nerve fibers, respectively. Scale bar = 10 μm.

**Supplementary Figure 4.**
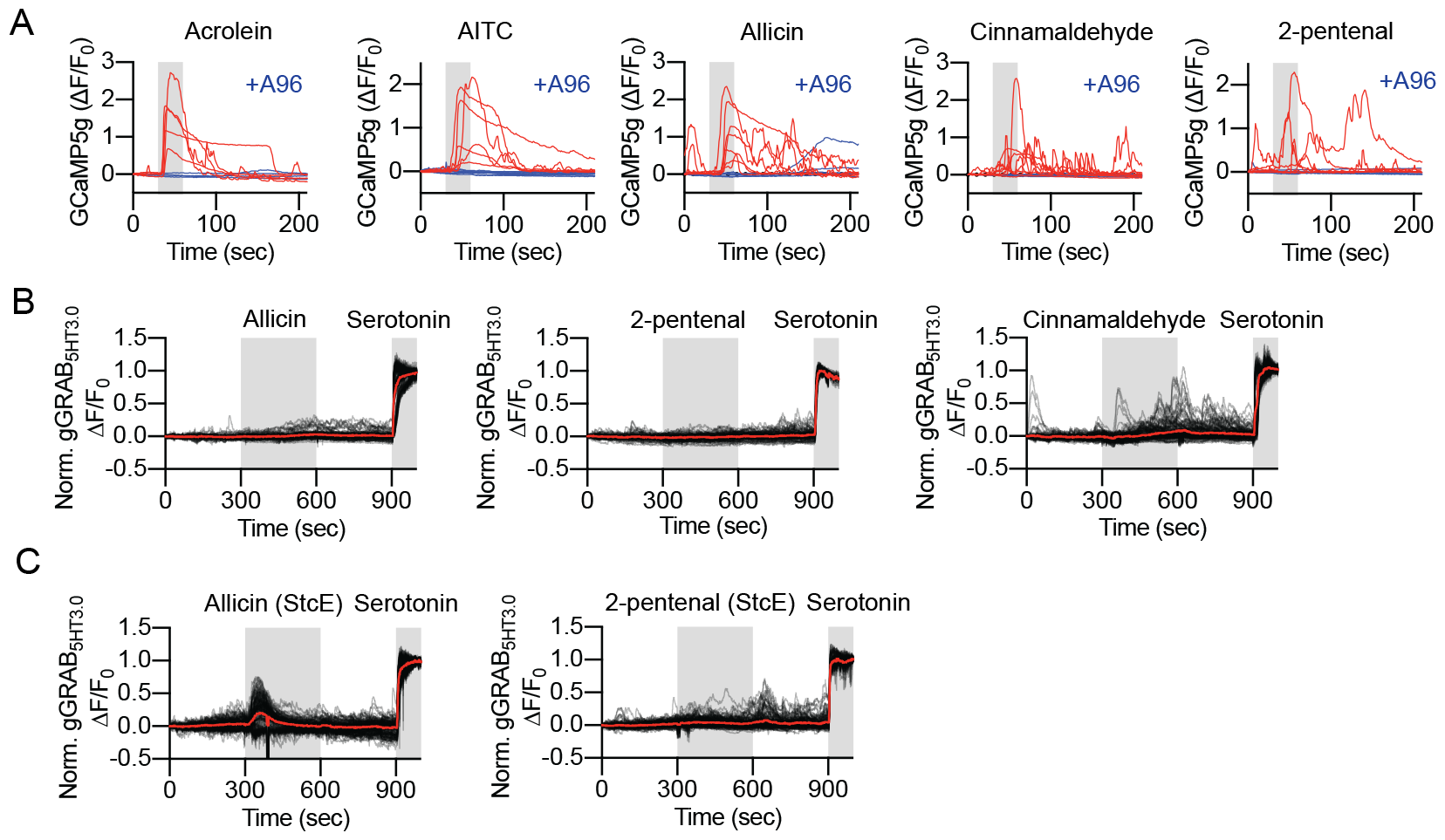
(A) Electrophiles (100 μM) activate EC cells in Tac1-IRES-Cre;GCaMP5g-IRES-tdTomato organoids. (B) Dietary electrophiles (100 μM) failed to activate crypt EC cells in the *ex vivo* prep isolated from Vil1-Cre;gGRAB_5HT3.0_ mice, with the exception of a minor yet statistically significant response elicited by cinnamaldehyde. (C) Electrophiles (100 μM) activated crypt EC cells after mucin digestion with StcE cells in the *ex vivo* prep isolated from Vil1-Cre;gGRAB_5HT3.0_ mice.

**Supplementary Figure 5.**
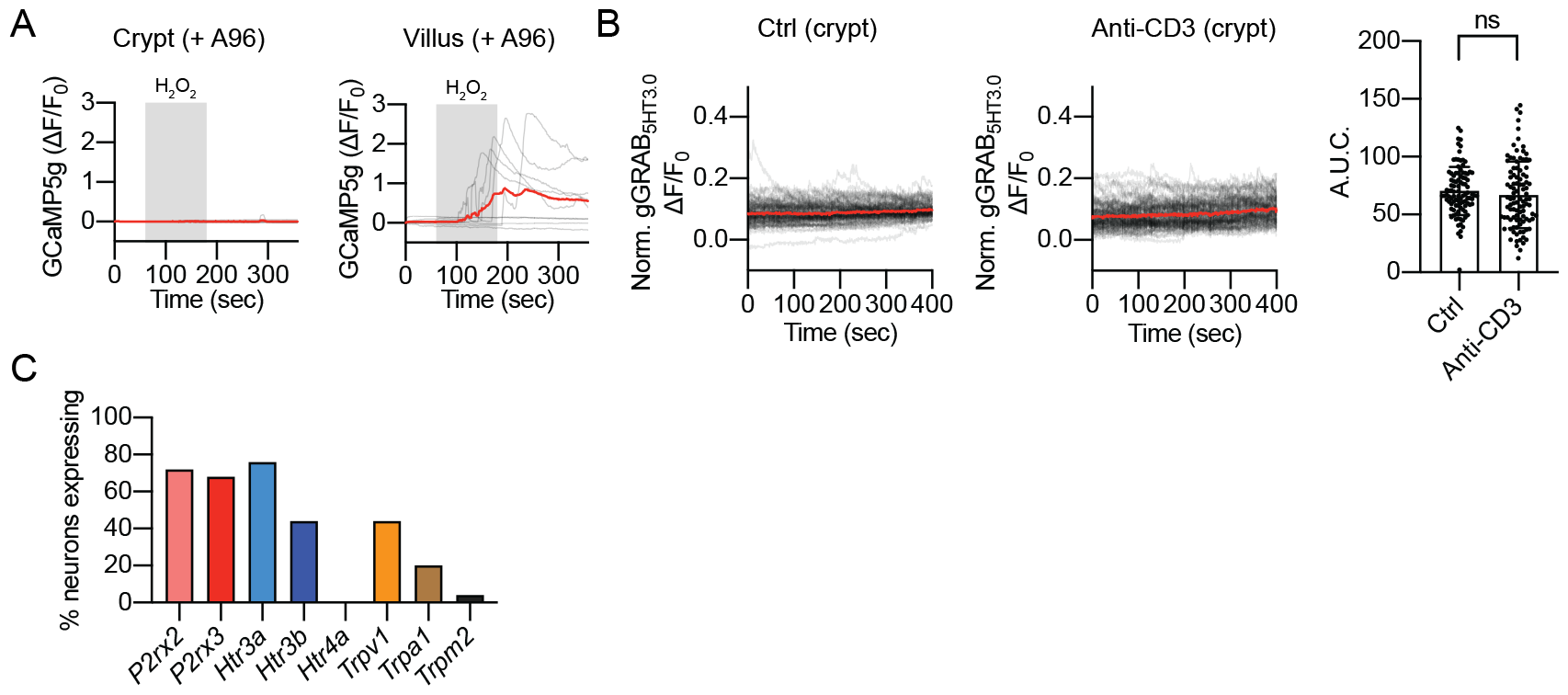
(A) Oxidative stress activates villus EC cells in a TRPA1-independent manner. Dissociated EC cells expressing GCaMP5g were exposed to 200 μM H_2_O_2_ for 2 minutes in the presence of 10 μM A96. (B) Oxidative stress does not activate crypt EC cells. The epithelial inflammation was induced by anti-CD3 antibody in gGRAB_5HT3.0_ sensor mice, and basal serotonin levels were measured in fleshly isolated *ex vivo* prep. The gGRAB_5HT3.0_ sensor was fully activated at the end of recordings for normalization. Welch’s t-test; ns: not significant. (C) Single-cell real-time PCR analysis of retrogradely traced small intestine mucosa-innervating vagal neurons. A target was defined to be present when a typical amplification curve was produced. n = 25

